# Early endosomes act as local exocytosis hubs to repair endothelial membrane damage

**DOI:** 10.1101/2022.11.02.514845

**Authors:** Nikita Raj, Lilo Greune, Martin Kahms, Karina Mildner, Rico Franzkoch, Olympia Ekaterini Psathaki, Thomas Zobel, Dagmar Zeuschner, Jürgen Klingauf, Volker Gerke

## Abstract

The plasma membrane of a cell is subject to stresses causing ruptures that must be repaired immediately to preserve membrane integrity and ensure cell survival. Yet, the spatio-temporal membrane dynamics at the wound site and the source of membrane required for wound repair are poorly understood. Here, we show that early endosomes, previously only known to function in the uptake of extracellular material and its endocytic transport, are involved in plasma membrane repair in human endothelial cells. Using live-cell imaging and correlative light and electron microscopy, we demonstrate that membrane injury triggers a previously unknown exocytosis of early endosomes that is induced by Ca^2+^ entering through the wound. This exocytosis is restricted to the vicinity of the wound site and mediated by the endosomal SNARE VAMP2, which is crucial for efficient membrane repair. Thus, the here identified Ca^2+^-evoked and localized exocytosis of early endosomes supplies the membrane material required for rapid resealing of a damaged plasma membrane, thereby providing the first line of defense against damage in mechanically challenged endothelial cells.

## Introduction

The plasma membrane acts as a selective barrier for a eukaryotic cell and ensures the maintenance of cellular homeostasis and integrity. It is subject to damage caused by frequently encountered challenges, such as mechanical forces resulting from shear stress, mechanical stretch or cell migration through confined space ^[1] [2]^. The resulting membrane wounds must be repaired to avoid cytosolic leakage which would otherwise lead to cell death and eventually cause tissue damage. Defects in plasma membrane repair have been linked to numerous disease pathologies, e.g. muscular dystrophies, diabetes, cardiac failure, and certain bacterial infections ^[3]^. Several cell-intrinsic mechanisms have been proposed to aid in membrane repair, including recruitment of intracellular vesicles to plug the wound ^[4]^, internalization of the damaged membrane ^[5]^, shedding of the wounded membrane by ESCRT proteins ^[6] [7] [8]^, and wound constriction supported by cytoskeletal forces ^[9]^. All of these processes are triggered by the entry of Ca^2+^ through the wound, which serves as the immediate danger signal indicating plasma membrane damage ^[10]^.

Following Ca^2+^ entry, an assortment of molecular regulators that aid in the various stages of repair, including Ca^2+^-binding proteins such as annexins and dysferlins as well as ESCRT proteins, have been identified ^[11]^. The different molecular players most likely act in a wounding-context dependent manner, either synergistically or exclusively ^[12]^. Yet, a sole requirement for all wound repair responses is the resupply of membrane to the damage site, irrespective of the players involved ^[13]^. Even though this is a vital criterion for the initial resealing of the wound to prevent further Ca^2+^ entry and cytosolic exchange ^[1]^, the source of membrane that is required for wound closure and how it is delivered to the wound site is controversial ^[3] [13] [14] [15] [16]^.

Earlier studies in the field have reported that intracellular vesicles are recruited to the wound site to form a membrane patch which then fuses to the edges of the injured plasma membrane to aid in wound closure ^[16] [17]^. However, this patch has only been observed in exceptionally large cells such as *Xenopus* oocytes ^[18]^, sea urchin eggs ^[19]^ and *Drosophila* early-stage embryos ^[20]^, which were subjected to large membrane ruptures and wherein specialized organelles such as cortical granules seem to be the source of the intracellular vesicles. These results are not applicable to the membrane repair responses in somatic cells due to the cell size and lack of cortical granules for the membrane supply. Previous reports on membrane repair in somatic cells suggested that lysosomal exocytosis is induced following membrane damage ^[21]^ and is required for the release of acid sphingomyelinase, which in turn promotes endocytosis of the wounded membrane ^[22]^. However, other studies employing pharmacological inhibition of lysosome exocytosis reported this process to be dispensable for plasma membrane resealing in somatic cells ^[23]^. Yet another pool of not precisely defined intracellular vesicles has been implicated to support membrane wound closure by lowering the membrane tension which is increased upon damage ^[4] [19] [24]^. Thus, so far no consensus has been reached on the still ambiguous sources of membrane required for resealing and recent studies have rather focused on the context-dependent molecular players ^[25] [26] [27]^ or the later stages of repair after a diffusion barrier has been re-established ^[28] [29]^. The ambiguity concerning the source of membrane for the repair process has been also augmented by the lack of studies on the spatio-temporal membrane dynamics that occur immediately after wounding in somatic cells, i.e. within the first 20-30 *s* that comprise the time interval for resealing ^[18]^.

To address this long-standing conundrum and investigate the immediate dynamics following membrane wounding in somatic cells, we chose to study membrane repair in endothelial cells. Vascular endothelial cells are subject to constant mechanical stress resulting from blood flow and interactions with circulating blood cells and as a consequence, frequently experience physiological wounds ^[30]^. Failure to repair these wounds and hence maintain endothelial cell and vascular integrity has been associated with diseases including bleeding disorders ^[31]^, diabetes ^[32]^, and recently also COVID-19 ^[33] [34]^. Thus, endothelial cells serve as an ideal model system to address fundamental aspects of the resealing process such as the identity of the membrane source that supplies material to the plasma membrane for immediate and efficient repair. Here we show that early endosomes represent this membrane source. We identify a previously unknown role for early endosomes, regarded classically as the organelles for endocytic uptake and transport, and show that they undergo a Ca^2+^-triggered exocytosis in response to membrane wounding. This specialized early endosome exocytosis is spatially restricted to the vicinity of the wound site, occurs immediately upon wounding, and supports the repair process through SNARE (VAMP2) mediated exocytosis. Thus, early endosomes, the classical organelle for endocytic uptake, can be repurposed to undergo a Ca^2+^-evoked exocytosis to aid in a cellular stress response such as plasma membrane repair and thereby prevent membrane damage induced cell death.

## Results

### Endosomal vesicles disappear near plasma membrane wound sites

Human umbilical vein endothelial cells (HUVEC) were injured by two-photon laser ablation and the reestablishment of membrane integrity was measured using the membrane-impermeable dye, FM4-64 (Figures S1 and S2) ^[35]^. This assay showed efficient membrane resealing in the presence of Ca^2+^ within 30 *s* of wound induction (Figure S1B) ^[35] [36] [37]^. Internal membrane compartments have been proposed to aid PM repair, e.g. by forming a membrane plug or patch beneath the wound in very large cells such as *Xenopus* oocytes ^[2]^. Therefore, we investigated the potential association of various intracellular compartments with endothelial PM repair by recording the dynamic behavior of an array of organelle markers upon laser injury in HUVEC. Interestingly, none of the intracellular organelles analysed were recruited directly to the wound site (assessed by the circular wound region of interest, ROI), arguing against the formation of a large intracellular membrane patch [18] (Figures S3 and S4). This is in line with other studies suggesting the lack of an intracellular plug or patch during wound resealing in somatic cells ^[38]^ [39].

However, we observed a striking disappearance of early endosome (EE), and to a somewhat lesser extent, late endosome/lysosome (LEL), associated proteins near the wound site immediately upon membrane damage (Figure 1A; Videos S1 and S2). Loading early endosomes with fluorescently labelled transferrin revealed that both endosome-associated proteins, such as GFP-2xFYVE, and the endosomal cargo, i.e., transferrin, disappeared in response to wounding indicating the loss of entire endosomes near the wound site (Figure S3G and Video S3). When Ca^2+^ was depleted from the extracellular environment, membrane resealing in HUVEC was impaired (Figure S1) and importantly, the disappearance of endosomal vesicles was also inhibited (Figure S5). This suggests that the endosomal disappearance is Ca^2+^ triggered and likely associated with membrane repair.

**Figure 1.**
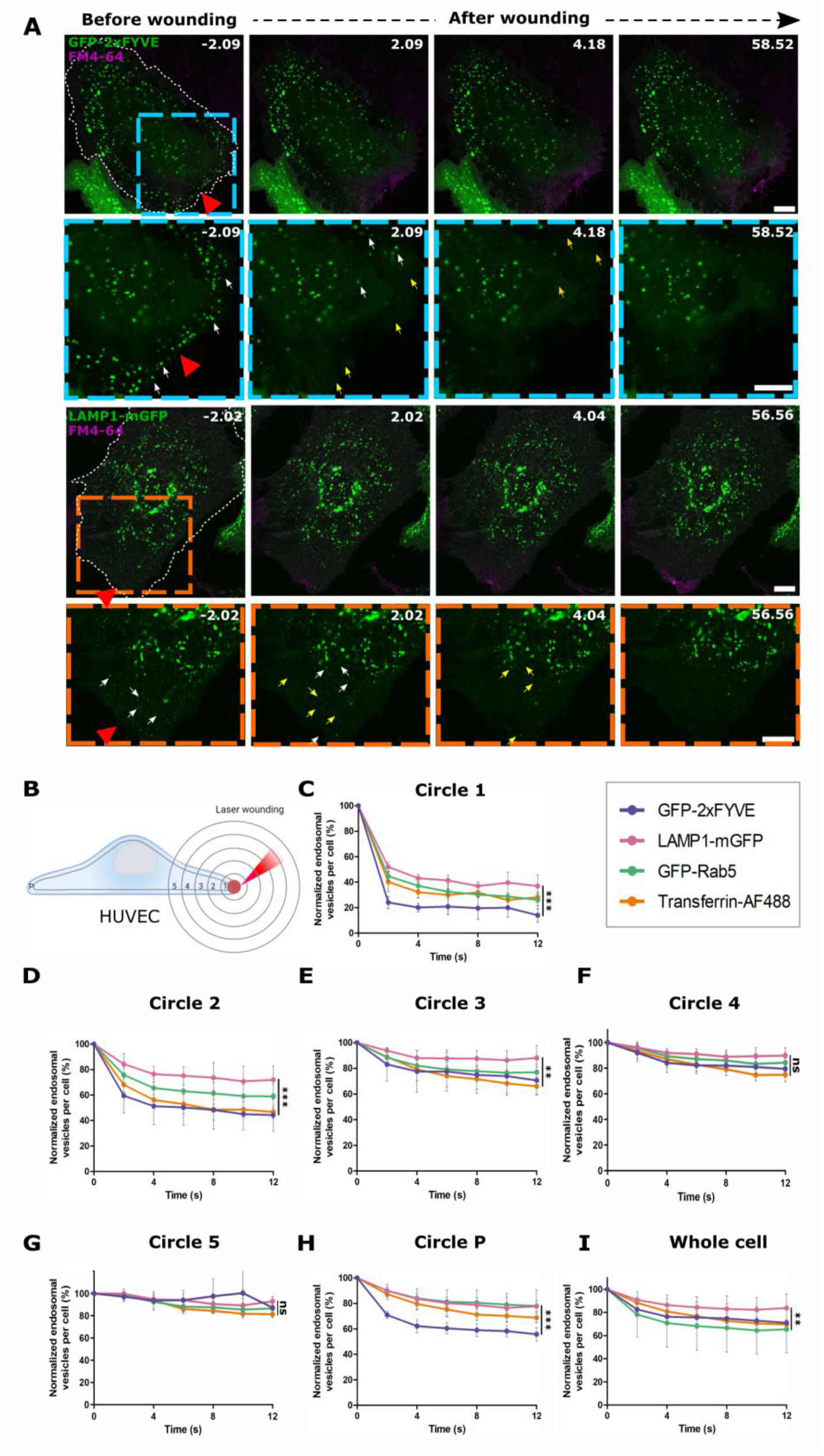
Membrane injury leads to the disappearance of early and late endosomal vesicles close to the wound site in HUVEC. (**A**) HUVEC transfected with GFP-2xFYVE or LAMP1-mGFP (green) and stained with the membrane dye FM4-64 (magenta) were subjected to laser injury while being imaged by time-lapse microscopy. Representative still images shown here. Red triangle, injury site. Boxed areas are magnified below; white arrows indicate endosomes before disappearance. Yellow arrows indicate the endosomal disappearance sites from the previous time frame. Wounded cells are outlined in white dashed lines. Scale bars, 10 μm; for zoom, 5 μm. (**B**) Scheme illustrating the ROI-analysis used to quantify disappearing endosomes with increasing distance from the wound site (red circle on the PM). (**C** to **H**) Quantification of the disappearance of endosomal markers over time, following laser injury in HUVEC transfected with GFP-2xFYVE (purple), GFP-Rab5 (green), LAMP1-mGFP (pink) or pulsed with transferrin-AF488 (orange), with t = 0 *s* representing the time of wounding. As in (B), endosomal disappearance is represented in circle 1 (C), circle 2 (D), circle 3 (E), circle 4 (F), circle 5 (G), circle P, the peripheral edge of the cell (H) and in the whole cell after wounding (I). Each data point shows the number of endosomes normalized to the initial number before wounding, expressed as a percentage, and mean + SD shown here. *n* = 18 - 21 cells, pooled from 3 independent experiments. The tests used for statistical comparison were: one-way ANOVA with Friedman test for (C to E) and (H), one-way ANOVA with Tukey’s test for (F and G), and one-way ANOVA with Kruskal-Wallis test (I). *ns*, not significant; \*\**P* < 0.01; \*\*\**P* < 0.001.

Employing a concentric ROI-based spatial segregation approach to quantify the wound-induced disappearances (Figure 1B), we next showed that the disappearance of endosomal proteins was restricted to the vicinity of the wound site, indicating a strictly localized response (Figures 1C-H). Moreover, it occurred within the first 10 s of wounding (Figures 1C-E) and always preceded resealing (Figure S1). Among the various endosomal proteins analysed, EE markers showed the greatest (~ 60 - 80 %) disappearance upon wounding (Figures 1C-H). By analyzing the behavior of endosomal markers in non-wounded cells, we further clarified that the endosomal disappearance was a wounding-specific response and not due to the diffusion of the endosomes over time and/or bleaching of the fluorescent endosomal markers by the ablation laser (Figure S6). Thus, EE and to a lesser extent LEL vesicles near the wound site disappeared immediately upon wound formation in HUVEC, indicating a possible role of these responses in plasma membrane repair.

### Late endosomal/lysosomal exocytosis is not required for PM repair in HUVEC

Although the role of lysosomes in PM repair is ambiguous ^[14]^, lysosomes have been implicated in aiding membrane repair in some cell types by undergoing exocytosis upon wound-induced Ca^2+^ elevation ^[40] [21]^. This prompted us to address whether the disappearance of LEL observed upon HUVEC wounding is due to their exocytosis. Therefore, we expressed the fusion construct pHluorin-LAMP1-mApple in HUVEC, which enabled the identification of the fusion of LAMP1-positive vesicles (i.e., LEL) with the PM as an increase in fluorescence of the pH-sensitive pHluorin ^[41] [42]^. We observed a disappearance in the LAMP1-mApple fluorescence, similar to Figure 1A, but also noted an increased fluorescence in the pHluorin channel at the site of disappearance, indicating exocytosis events (Figure S7A). The validity of the pHluorin-LAMP1-mApple fusion construct in recording LEL exocytosis was also confirmed without wounding but by using a conventional Ca^2+^ ionophore, ionomycin, which has been shown to induce Ca^2+^ elevation and thereby LEL exocytosis ^[43]^ (Figure S7B). Taken together, we conclude that the LEL disappearances observed immediately upon wounding in HUVEC reflect exocytosis events, which have been linked to membrane repair in other cells ^[21]^.

Next, we investigated whether the wounding-induced LEL exocytosis is functionally involved in HUVEC membrane repair by applying a well-characterized inhibitor of lysosomal exocytosis, U18666A ^[44] [45]^. As expected, U18666A caused a marked inhibition of LEL exocytosis upon wounding (Figures 2A and 2B) and ionomycin treatment (Figure 2C and S8A), respectively. However, upon wounding of HUVEC treated with U18666A, no defect in membrane resealing was observed (Figures 2D and 2E). This result was confirmed using different mechanical membrane wounding techniques - cell scraping (Figure 2H and S9B) and glass beads mediated injury (Figure 2I). These population wounding assays reflect an end point of membrane repair rather than the real-time kinetics recorded in the laser wounding assay ^[46]^, and thus serve as complimentary methods to study the involvement of lysosomal exocytosis. This also accounts for the different resealing efficiencies in the controls (in Figures 2H and 2I) as observed before in different cells ^[47] [48]^, which are enhanced by the susceptibility of the flat endothelial cells (Figure S2B) to detachment via scrape injury or via glass wounding (Figures S9B and S9C) ^[49] [50]^. However, albeit these technical limitations in HUVEC, all the wounding methods applied here indicated that blocking lysosomal exocytosis is not required for endothelial membrane repair.

**Figure 2.**
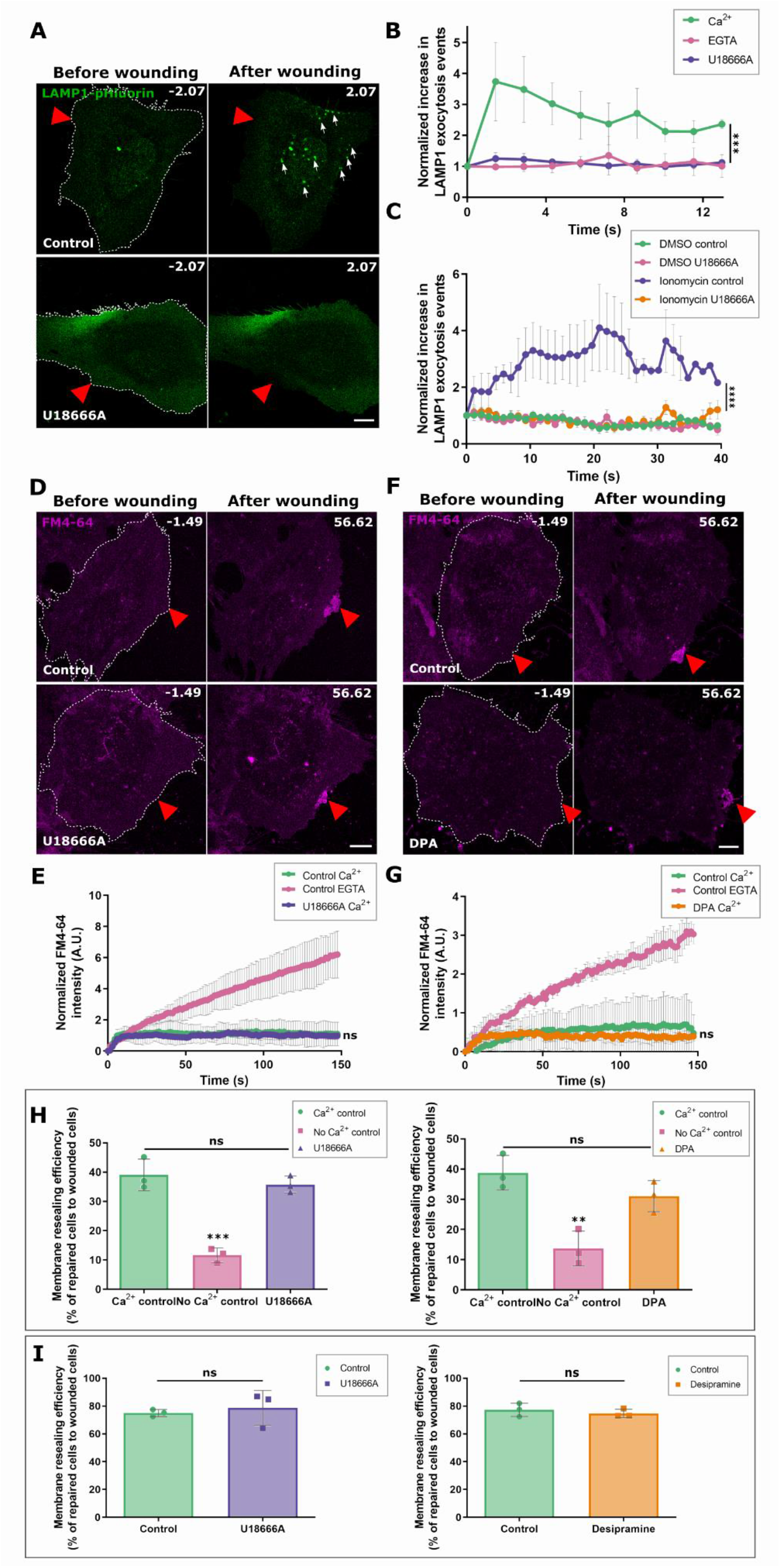
LEL exocytosis does not contribute to HUVEC membrane repair. **(A** and **B)** pHluorin-LAMP1-mApple expressing HUVEC were laser wounded in the presence of Ca^2+^ (+ U18666A treatment) or EGTA. Still images of LAMP1-pHluorin (green) are shown here. White arrows highlight LEL (LAMP1) exocytosis events. LAMP1-exocytosis events normalized to the initial count before wounding, plotted over time. (**C**) Ionomycin induced LAMP1-exocytosis events (see Figure S8A) were quantified over time in the absence or presence of U18666A and normalized to the initial punctae count before ionomycin addition. (**D and E**) Still images of laser injury of HUVEC treated with U18666A in the presence of FM4-64 (magenta), and kinetics of membrane resealing quantified over time. (**F** and **G**) Laser injury of HUVEC treated with DPA in the presence of FM4-64 with resealing dynamics quantified in G. (**H** and **I**) Resealing efficiency of cells treated with U18666A (purple) or DPA (orange) after mechanical scrape injury (H) or glass bead injury (I), represented as percentages. Red triangles, wound ROI. Mean + SEM plotted for (C) and mean + SD plotted for all other graphs. *n* = 22 – 25 cells (B and C), 24 - 26 cells (E), 24 – 28 cells (G), pooled from 3 independent experiments. *P* value is calculated using one-way ANOVA with Kruskal-Wallis test for (B) and (C), one way ANOVA with Hol-Sidak’s test (E), ordinary one-way ANOVA with Tukey’s test (G and H), and unpaired two-tailed Student’s *t*-test (I). *ns*, not significant; \*\**P* < 0.01; \*\*\**P* < 0.001; \*\*\*\**P* < 0.0001.

Previous work has shown that acid sphingomyelinase (ASM) is released during wound-induced lysosomal exocytosis and that the secreted ASM is responsible for supporting an endocytosis-mediated PM repair ^[22] [51] [52]^. As our data revealed that LEL exocytosis is not required for PM resealing in HUVEC, we next analysed whether ASM, possibly released from sources other than LEL, regulates HUVEC membrane repair. Therefore, we employed desipramine (DPA) which has been shown to block ASM function ^[53] [54]^. Upon inhibition of ASM activity by DPA in HUVEC (Figure S8B), membrane resealing was unperturbed following laser wounding (Figures 2F and 2G) and in the different mechanical wounding assays (Figures 2H, 2I and S9C). Together, these findings show that lysosomal exocytosis and the release of ASM are not required for endothelial PM repair.

### Spatially confined early endosome exocytosis occurs upon PM injury

As we ruled out the functional involvement of LEL in HUVEC membrane resealing, we next examined the role of EE, which disappeared near the wound site in considerably higher numbers as compared to LEL (Figures 1C-E). We first verified that other EE-associated proteins, expressed as GFP fusions in HUVEC, disappeared near the wound site in HUVEC, similar to GFP-2xFYVE shown in Figure 1A, indicating a general response of EE to membrane damage (Figure S10). We next assessed whether the EE disappearances were due to exocytosis events, owing to the LEL disappearances indicating LEL exocytosis. Several studies have described the fusion of recycling vesicles with the PM and also the homotypic fusion of EE, but to date, very little is known about EE exocytosis, in particular, evoked EE exocytosis that had not been observed before ^[55] [56]^. To analyse whether the EE disappearances reflect exocytosis events, we utilized an EE membrane protein, transferrin receptor (TfR), fused to a pH-sensitive reporter, TfR-pHuji ^[57]^. Laser injury of HUVEC expressing the EE marker, GFP-2xFYVE, together with TfR-pHuji showed the disappearance of 2xFYVE vesicles, as observed before, and interestingly a concomitant increase in TfR-pHuji fluorescence at the sites of 2xFYVE disappearance (Figure 3A; Video S4). The increased TfR-pHuji fluorescence intensity at the cell surface often appeared in clusters detected close to the wound site and immediately after membrane damage (Figure 3B and S11A-B). A significant fraction of these TfR clusters near the wound site overlapped with the sites of GFP-2xFYVE disappearance, strongly suggesting the occurrence of evoked exocytotic fusion events of EE which had not been observed before (Figures 3C and S11C).

**Figure 3.**
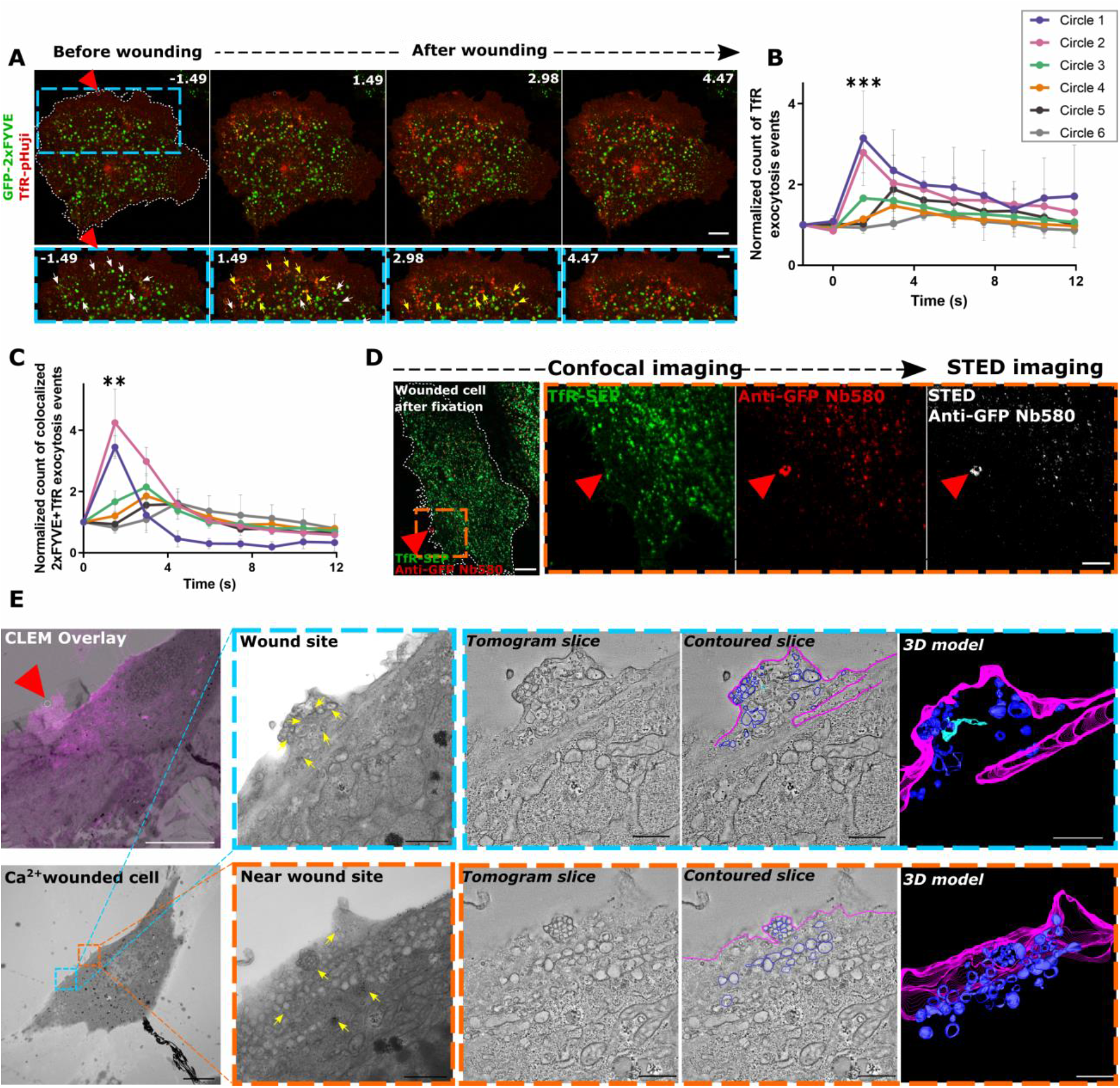
EE preferentially undergo exocytosis near the wound site. (**A**) HUVEC transfected with GFP-2xFYVE (green) and TfR-pHuji (red) were laser-wounded. Blue dashed boxes of each time point are magnified below; white arrows indicate 2xFYVE endosomes before disappearance; yellow arrows indicate TfR exocytosis events at the sites of 2xFYVE disappearance (**B**) The increase in TfR exocytosis events (clusters with increased pHuji signal) was quantified over time, represented in zones of increasing distance from the wound site (see Figure S11A), and normalized to the baseline count in each cell per μm^2^ of ROI area. (**C**) EE exocytosis events positive for concomitant 2xFYVE disappearance and TfR exocytotic cluster formation (see A), quantified as in (B) over time with each ROI area and initial count normalized. (**D**) Detection of TfR exocytosis events around the wound site (orange box magnified on the right) by confocal and STED imaging, following the anti-GFP nanobody exocytosis assay. Image representative of 4 experiments. (**E**) HUVEC were labelled with BSA-gold, washed, laser wounded in the presence of FM4-64, fixed, and processed for TEM. CLEM overlay image of the relocated wounded cell (0-250 nm section). Conventional TEM image of the consecutive 250 nm section (from 250-500 nm height) shown below with the wound site marked by a blue box and a nearby site marked by an orange box. Magnifications are shown on the top right for the wound site and the bottom right for the nearby site. Shown next to each TEM zoom-in is a representative tomographic slice extracted from each double tilt tomography series followed by the contoured slice and the 3D model of the entire tomogram (see Videos 6 and 7). In the contoured slice and the model, PM (magenta) and unambiguously identified endosomes, which are vesicular (dark blue) and tubular (light blue), are traced. Yellow arrows indicate BSA-gold labelled membranes. Representative CLEM image from *n* = 17 cells over 3 experiments. Red triangles, wound sites. Scale bars, 10 μm (A, D, and E); 1 μm in magnified TEM views and 500 nm in tomographic slices of (E)]. *n* = 22 - 29 cells for (B) and (C), from 3 independent experiments. Statistical comparisons were performed with ordinary one-way ANOVA for (B) and one-way ANOVA with Kruskal-Wallis test for (C). \*\**P* < 0.01; \*\*\**P* < 0.001.

To further confirm that the wounding-induced TfR surface clusters correspond to EE exocytosis, we adapted an exocytosis detection protocol ^[58]^. We expressed TfR-SEP (GFP variant of TfR-pHuji) in HUVEC and blocked the resident surface pool of TfR-SEP with an unconjugated anti-GFP nanobody. Thereafter, new TfR exocytosis events could be imaged by adding fluorescently labelled anti-GFP nanobodies during laser injury. Confocal and STED imaging of the laser-injured cells fixed immediately after wounding showed an accumulation of TfR positive exocytosis events at and around the wound site but not further away from the wound site (Figures 3D and S12). The TfR exocytosis events right at or very close to the wound site were not observed during live confocal imaging as noted by a lack of an appearance of TfR-pHuji at the wound site (see Figure 3A and TfR-SEP panel of Figures 3D and S12A). This indicates that the nanobody labelling protocol permits the capture of very transient exocytosis events occurring possibly at the edges of the wound site, which can then be observed in the fixed cells.

In addition, ultrastructural analysis of the HUVEC wound sites by correlative light and electron microscopy (CLEM) (Figure S13) showed an abundance of convoluted vesicular membranes, likely of endosomal origin as revealed by internalized BSA-gold labelling (Figure 3E blue inset; Video S6). This enrichment around the edges of the wound site of the fixed CLEM image is consistent with the nanobody labelled STED images representing a transient snapshot of the membrane repair process. An extensive accumulation of vesicular structures, again most likely of endosomal origin, was also observed near the wound site, similar to the TfR exocytotic events noted earlier (Figure 3E orange inset; Video S7). This vesicle accumulation was not seen in a region farther away from the wound site nor in a neighbouring unwounded cell or a cell wounded in the absence of Ca^2+^ (Figures S14-S15 and Videos S8 and S9, respectively). Collectively, these experiments indicate that early endosomes undergo wound-induced exocytosis, which occurs immediately after wounding and is spatially localized to the vicinity of the injury site.

The possibility that such spatially restricted EE exocytosis could occur in response to membrane damage was probably neglected so far because homotypic EE fusions and endosomal recycling are not known to be Ca^2+^ dependent ^[1]^. However, as our results suggested that the EE disappearance is driven by Ca^2+^ entering the cell through the wound, we next analysed in detail the interdependence of wound-induced Ca^2+^ dynamics and EE exocytosis. Along with the dependence of the observed EE exocytosis on extracellular Ca^2+^ (Figures 4A and S5), the EE disappearances also followed the dynamics of an injury-evoked Ca^2+^ wave, which was tracked using the Ca^2+^ indicator R-GECO 1.2 (Figures 4B, 4C and S11D). Thus, Ca^2+^ entering through the wound most likely triggered the exocytosis of EE residing in proximity to the wound site. Interestingly, physiological stimuli known to elicit cytosolic Ca^2+^ elevation such as histamine did not induce the exocytosis of EE, most likely because they failed to produce the high Ca^2+^ concentration encountered close to the wound site upon injury (Figure 4D and Video S5). Thus, membrane damage triggers a Ca^2+^-induced exocytosis of EE that most likely supports instantaneous PM resealing.

**Figure 4.**
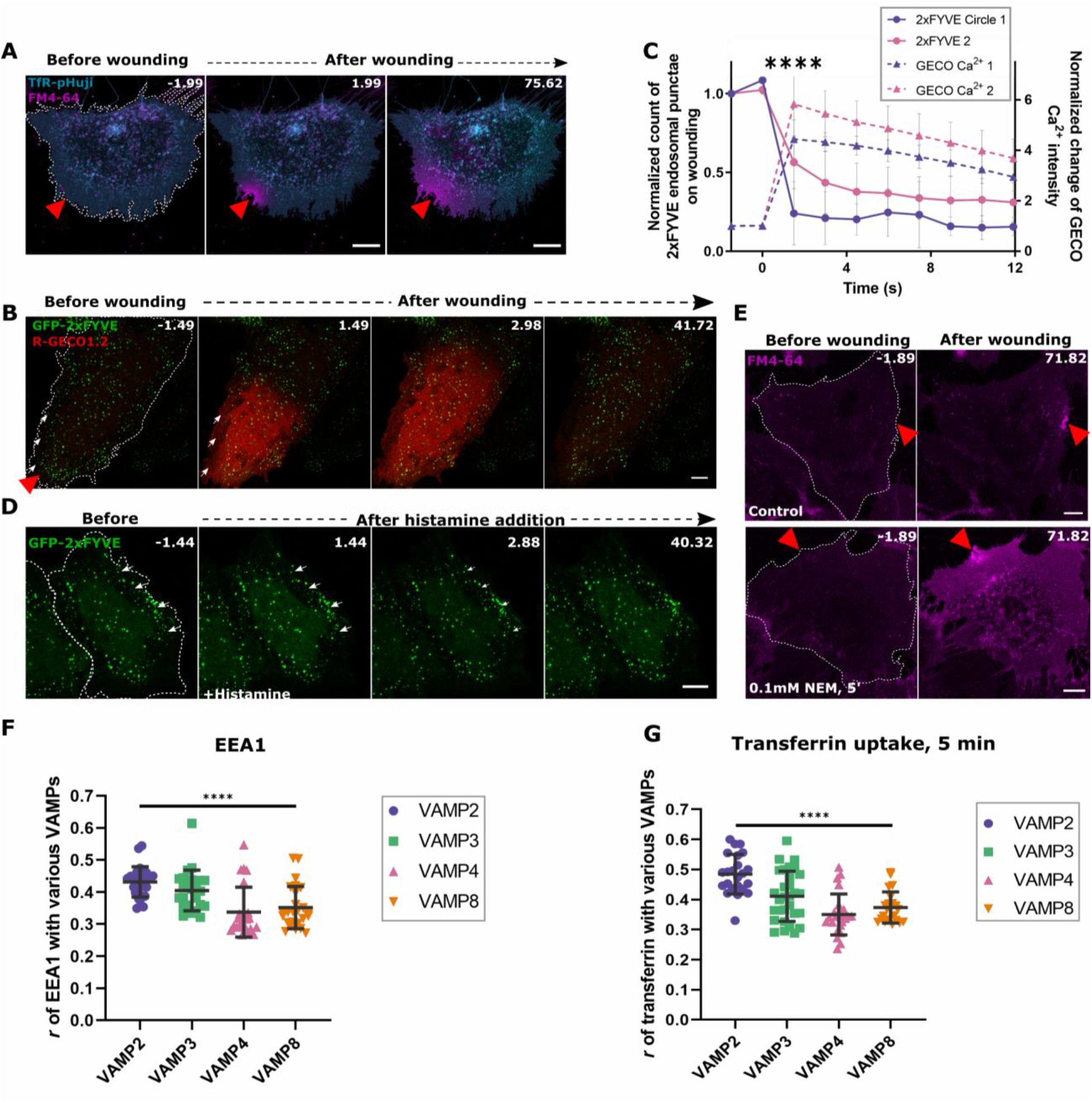
Exocytosis of early endosomes occurs solely in the presence of high levels of Ca2+ and is dependent on SNARE-mediated fusions. **(A)** TfR-pHuji expressing HUVEC (displayed in cyan) were laser wounded in the presence of EGTA and the wounding response was recorded using FM4-64 (magenta). Representative images before and after wounding are shown. No change in the localization of TfR-pHuji fluorescence was observed. (**B**) HUVEC expressing GFP-2xFYVE (green) and the Ca^2+^ indicator, R-GECO 1.2 (red) were laser wounded. Images show Ca^2+^ entry and white arrows indicate disappearing 2xFYVE endosomes in the corresponding frames. (**C**) Overlap of GFP-2xFYVE disappearance (solid lines) with the GECO Ca^2+^ wave intensity increase (dashed lines) quantified for circles 1 and 2. (**D**) GFP-2xFYVE expressing HUVEC were recorded live and 100 μM histamine was then added to induce intracellular Ca^2+^ elevation. Time-lapse images show no change in the overall number of GFP-2xFYVE positive endosomes and a few such examples are marked by white arrows. **(E)** HUVEC were treated with the SNARE inhibitor, NEM (0.1 mM, 5min), or vehicle control and subjected to laser ablation in the presence of FM4-64 (magenta). Images pre and post wounding are shown. **(F** and **G)** Colocalization of the various VAMP proteins with respect to the EE marker, EEA1 (F) and transferrin-AF647 labelled for 5 min (G), was calculated using Pearson’s correlation coefficient, *r*, and represented as distribution plots with mean + SD. Red triangles, wound sites and white dashes indicate the laser ablated (A, B, E) and histamine stimulated (D) cells. Scale bars, 10 μm and images representative of 4 independent experiments (A, D, and E). *n* = 22 - 29 cells for (C) and 25-30 cells for (F and G), from 3 independent experiments. Statistical comparisons were performed using two-tailed Mann-Whitney *U* test (C), and one-way ANOVA with Kruskal-Wallis test (F and G), with *P* < 0.0001.

### VAMP2 promotes early endosomal exocytosis during HUVEC resealing

We next aimed to investigate how EE undergo exocytosis during membrane repair in HUVEC. As Ca^2+^ regulated EE exocytosis had not been described before, we analysed the role of classical exocytotic fusion proteins, SNAREs ^[59]^. Inhibition of SNARE mediated fusion by NEM (*N*-ethylmaleimide) ^[60]^ showed an immediate inhibitory effect on HUVEC membrane resealing (Figure 4E) indicating a role for SNARE mediated membrane fusions in PM repair and most likely an underlying wound-induced EE exocytosis. Of the endosomal v-SNAREs analysed in HUVEC, VAMP2 and to a lesser extent VAMP3, were unambiguously identified on EE. These EE populations were also positive for the FYVE domain-containing early endosome-associated protein 1 as well as internalized transferrin (Figures 4F, 4G and S16). Intriguingly, VAMP2-SEP expressed in HUVEC, upon wounding formed cluster-like structures of increased fluorescence close to the wound site, comparable to the TfR exocytotic clusters observed earlier (Figures 5A and 5B; Video S10). In contrast, hardly any wound-induced exocytotic clusters were seen for VAMP3, VAMP4, and VAMP8 (Figures S17-S18). The VAMP2 positive SEP clusters were thus indicative of specific SNARE mediated exocytotic events that occur during PM resealing.

**Figure 5.**
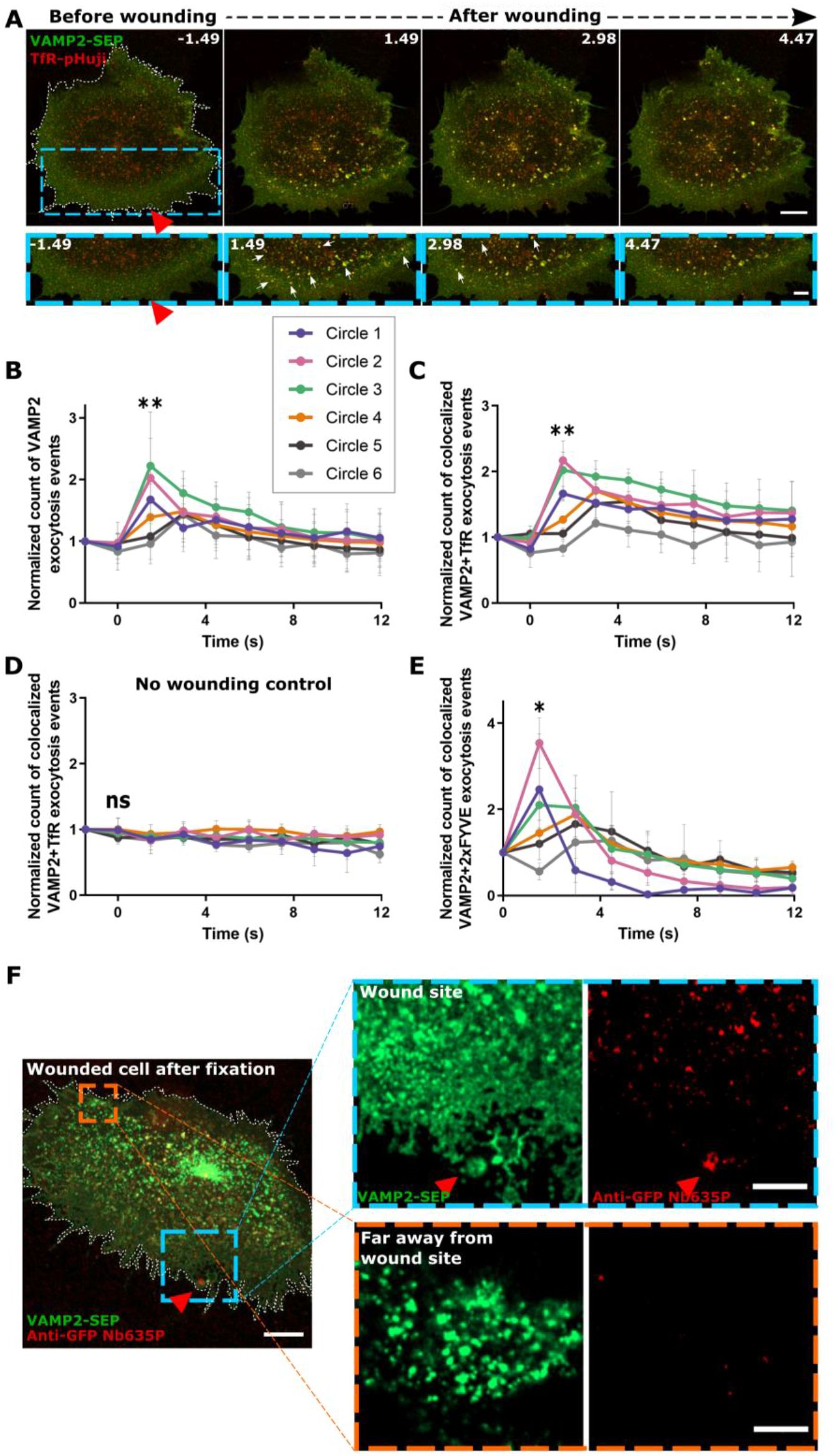
VAMP2 participates in wound induced EE exocytosis. **(A)** HUVEC cotransfected with VAMP2-SEP (green) and TfR-pHuji (red) were laser wounded and sequential images are shown. Blue dashed boxes are magnified below; white arrows indicate VAMP2 exocytotic events at the sites of TfR exocytosis. (**B**) Quantification of exocytosis events positive for VAMP2-SEP fluorescence increase over time, in regions of increasing distance from the wound site (Figure S11A), normalized to the circle ROI area and punctae count before wounding. (**C**) EE exocytosis events positive for colocalized VAMP2-SEP and TfR-pHuji signals of increased intensity, quantified as in (B), and normalized to the area of each ROI and baseline colocalized punctae count. (**D**) Similar analysis as in (C) for cells subjected to low laser power ablation (non-wounded control). (**E**) Analysis of exocytotic events showing VAMP2-SEP fluorescence increase that colocalized with mApple-2xFYVE disappearance, plotted over time. (**F**) Detection of VAMP2-SEP exocytosis events around the wound site (magnified in blue) and far away from the wound site (orange box) by the anti-GFP nanobody exocytosis assay. Red triangle, wound site. Scale bars, 10 μm; 5 μm in zooms (for A and F). *n* = 21 cells (C), *n* = 23 (D) and *n* = 19 (E), from 3 independent experiments, and for (B), *n* = 39 cells from 6 experiments. Statistical comparisons were performed using one-way ANOVA with Kruskal-Wallis test for (B), (C), and (E), and for (D), one-way ANOVA was used. *ns*, not significant; \**P* < 0.05; \*\**P* < 0.01.

We next explored whether the VAMP mediated fusions correspond to EE exocytosis events in HUVEC, by wounding cells expressing TfR-pHuji together with various VAMP constructs. VAMP2-positive clusters, especially those appearing close to the wound site, strongly colocalized with the wound induced TfR exocytotic events (Figures 5A, C, D; Video S10). Clusters of increased VAMP2-SEP fluorescence, which showed maximal association to TfR exocytotic clusters, also corresponded to the sites of EE (mApple-2xFYVE) disappearance, as seen with TfR (Figures 5E, S19 and S20G). Analysis of VAMP3, VAMP4, and VAMP8 revealed very few exocytosis events that barely overlapped with EE fusions (Figures S20A-F). Finally, using the anti-GFP nanobody-based exocytosis detection assay, we reaffirmed that the SEP-positive VAMP2 clusters indeed correspond to exocytosis events that occur transiently at and predominantly around the wound site upon membrane damage (Figure 5F). Together, these results suggest that VAMP2 is involved in regulating this previously unrecognized Ca^2+^-evoked EE exocytosis events which occurs in response to HUVEC membrane damage.

### VAMP2 mediated early endosome exocytosis is required for HUVEC resealing

The identification of VAMP2 as a SNARE associated with EE that undergo wound-induced exocytosis prompted us to probe for its functional role in HUVEC membrane repair. siRNA mediated depletion of VAMP2 in HUVEC (Figure 6A) resulted in significant impairment of HUVEC membrane resealing, as revealed by the drastic uptake of FM4-64 dye after laser injury (Figures 6B and 6C). We also probed cells depleted of VAMP3 (Figure S21A) as a control for a SNARE that did not show a prominent association with wounding-induced exocytotic fusions of EE (Figures S16 and S17). Here, no defects in membrane resealing were observed, arguing for an EE SNARE (VAMP2)-specific involvement in membrane repair (Figure 21B). These findings were corroborated using an EmGFP-based miRNA system to knock down VAMP2 protein levels in cells identifiable by their co-cistronic EmGFP expression (Figure 6D). Again, HUVEC with miRNA mediated depletion of VAMP2 showed a severe membrane resealing defect (Figures 6E and 6F). The strong repair defect observed here in individual cells with maximal VAMP2-depletion, selected based on the highest GFP expression, even surpasses the defect in Ca^2+^-free control cells. This is most likely due to the abundant binding of FM4-64 dye to intracellular membranes and complexes that remain exposed in the non-resealed cell and fail to fuse with the PM in the presence of Ca^2+^ owing to loss of VAMP2. Similarly, a marked inhibition of membrane resealing was observed in VAMP2-depleted HUVEC subjected to mechanical scrape and glass bead membrane wounding (Figures 6G, 6H and S22). The differences in the extent of repair efficiency in the VAMP2-depleted cells between laser wounding and mechanical wounding population assays is probably due to the siRNA vs GFP-based miRNA systems used ^[61]^. The miRNA system allowed to specifically select single cells with maximal depletion while the siRNA VAMP2 transfection results in a population of cells with varying levels of VAMP2 knockdown. Using the EmGFP-miRNA system, we also confirmed that VAMP2 mediated EE exocytosis is necessary for HUVEC membrane repair, as TfR exocytotic events around the wound site were significantly reduced upon membrane damage in VAMP2-depleted cells (Figure 6I). Thus, VAMP2 mediated EE exocytosis is crucial for membrane repair in HUVEC, most likely by rapidly supplying extra membrane to be used in the PM resealing process (Figure 6J).

**Figure 6.**
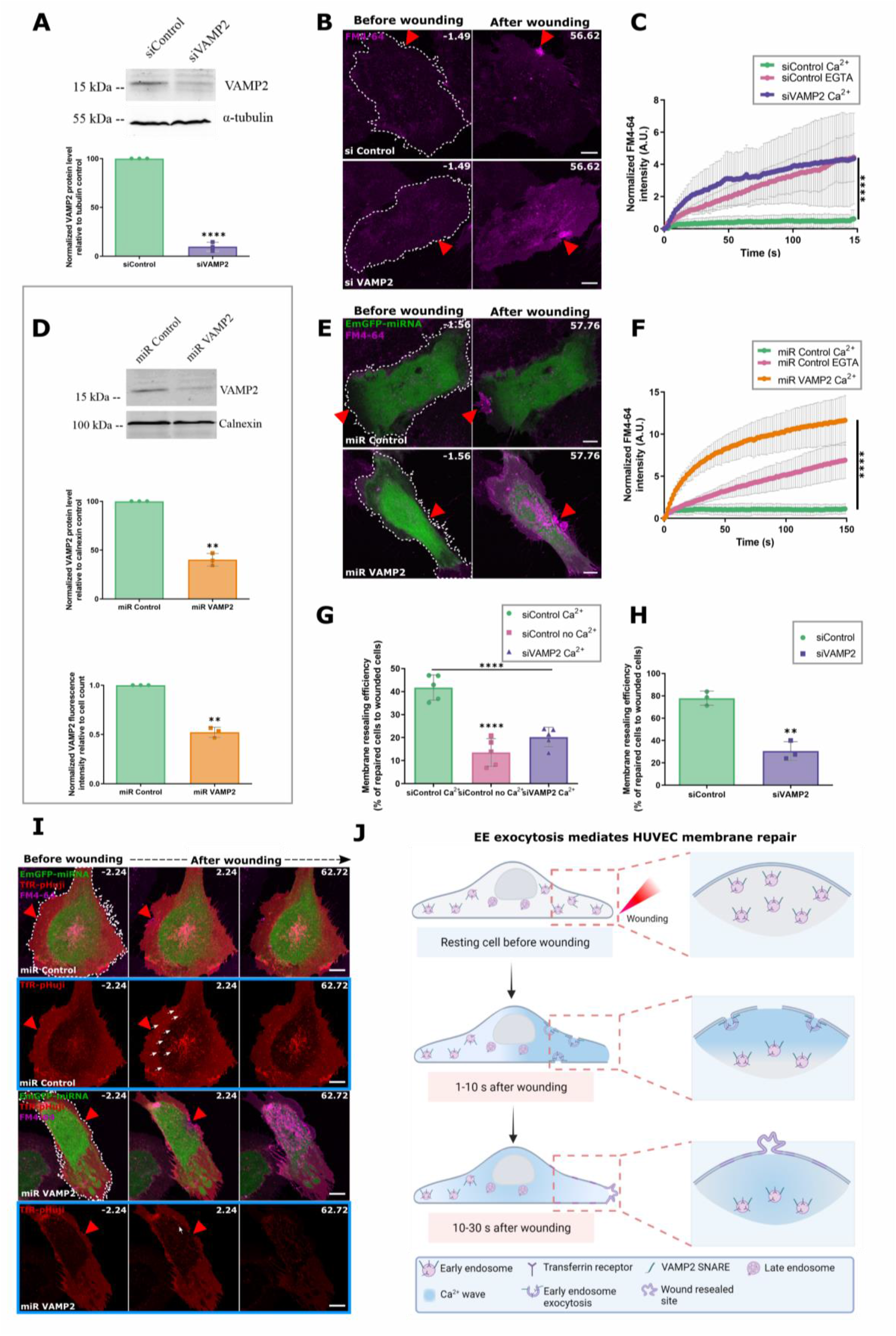
EE exocytosis regulated by VAMP2 is critical for HUVEC membrane resealing. (**A**) Immunoblot showing levels of VAMP2 after siRNA treatment with siControl (siC) or siVAMP2 pool (top panel) and represented as a percentage to the loading control tubulin (bottom). (**B** and **C**) siC or pooled siVAMP2 transfected HUVEC in the presence of FM4-64 (magenta) were laser-wounded (B) and the resealing kinetics quantified (C). (**D**) HUVEC were transfected with Em-GFP miRNA control (miC) or miRNA VAMP2 and assessed for VAMP2 knockdown by western blot of cell lysates (VAMP2 levels are normalized to loading control, calnexin) and flow cytometry analysis of VAMP2 staining (bottom panel). **(E** and **F**) EmGFP-miRNA control or miRNA VAMP2 transfected cells were laser wounded and resealing analyzed by FM4-64 dye influx (E) and quantified (F). (**G** and **H**) siC or siVAMP2 pool transfected cells were analyzed for membrane resealing after mechanical scrape injury (G) or glass bead injury (H). (**I**) VAMP2 depletion interferes with wounding induced exocytosis of TfR positive EE. HUVEC were transfected with Em-GFP miC or miR VAMP2 together with TfR-pHuji (red) and laser wounded in the presence of FM4-64 dye. The TfR channel is highlighted below in blue boxes, and white arrows indicate EE exocytosis events positive for TfR-pHuji fluorescence. (J) Model of the wounding induced exocytosis of EE for wound repair. Red triangles, wound sites.Mean + SD plotted for all graphs. *n* = 24 - 26 cells, pooled from 3 independent experiments in (C) and (F). Tests used for comparisons: unpaired two-tailed Student’s *t*-test ( A),(D), and (H), one-way ANOVA with Kruskal-Wallis test for (C) and (F), and one-way ANOVA with Tukey’s test (G). \*\**P* < 0.01; \*\*\**P* < 0.001; \*\*\*\**P* < 0.0001.

## Discussion

Our results identify a novel role for early endosomes, which are conventionally known to participate in endocytic transport, in plasma membrane repair of human endothelial cells. EE close to the wound site rapidly fuse with the PM and this process is required for efficient repair most likely by providing additional membrane for the resealing. Previously, lysosomes have been considered as the predominant source of membrane needed for wound resealing based mainly on their well-documented exocytotic response to increased cytosolic Ca^2+^, a hallmark of wounded cells ^[42] [40] [62] [63] [21]^. For the same reason, early endosomes had been excluded as they typically do not undergo exocytosis in response to physiologically encountered Ca^2+^ levels. We now show that early endosomes can indeed undergo exocytosis in response to Ca^2+^, however, only at the exceedingly high Ca^2+^ concentrations met in the vicinity of the wound site in injured cells.

The strict dependence of EE exocytosis on elevated Ca^2+^ levels is substantially different from the behavior of EE in resting or agonist-activated endothelial cells, suggesting a specific repurposing of EE in the danger situation of membrane disruption. The abundant and fast exocytotic response of EE in HUVEC (50-70% of EE around the wound site fuse with the PM) is likely to be necessary for immediate resealing of wounds as opposed to the much lower levels of lysosomal exocytosis observed in wounded HUVEC (correlating to the lack of an abundant pool of peripheral lysosomes) ^[42] [40]^. Moreover, in contrast to the VAMP2-mediated exocytosis of EE, which is functionally relevant for PM repair, lysosomal exocytosis is not required to support membrane resealing in HUVEC, but is rather shown to be a Ca^2+^ mediated response initiated by wound induced Ca^2+^-influx. This is in line with other studies, which have demonstrated unperturbed membrane resealing after blocking lysosomal exocytosis using vacuolin, suggesting a vesicle pool other than lysosomes as the membrane source for wound resealing ^[23]^.

Interestingly, most studies showing the involvement of lysosomes focus on wounds generated using pore-forming toxins, which are much smaller (<100 nm in diameter) ^[13] [50]^ than wounds generated *in vivo* due to mechanical stresses ^[64]^, suggesting that different repair strategies can operate depending on wound size. In the case of the larger, mechanically inflicted wounds, EE are ideally suited to provide additional membrane as they reside in the vicinity of the PM and thus can respond rapidly to wounding without requiring extensive transport to the wound site.

Evolutionary modifications of Ca^2+^ regulated exocytosis have been long speculated to mediate PM remodeling in different contexts ^[19]^. We propose that EE exocytosis in HUVEC is a specialized Ca^2+^ triggered, emergency-based process that occurs upon wound formation. This is of particular importance in endothelial cells, which are frequently subjected to wounds ^[30]^ and require fast and robust membrane delivery to support PM resealing. It is plausible that cells utilize early endosomes in such conditions, not just due to their abundance and proximity to the membrane, but also because they are easily replenished via ongoing uptake of PM by endocytosis. Endothelial cells are likely to respond to PM injury by rapid membrane delivery as they are very flat, and the ample peripheral EE are close to the PM in any given region of the cell. Moreover, these cells must have developed an efficient PM repair machinery to prevent wound-induced endothelial leakage and a resulting inflammatory reaction in a constantly challenging environment ^[31] [65]^.

## Materials and methods

### Cell culture

Primary HUVEC (Human Umbilical cord Vein Endothelial Cells) were obtained from Promocell (C-12203) and were used either directly or cryopreserved for later use. Cells were cultured on Corning CellBind or collagen-coated dishes in an equal mixture of ECGM-2 (Promocell, C-22111) and M-199 (Pan Biotech, P04-07500), each supplemented with serum (10 % FCS for M199 (Capricorn scientific, FBS-11A)), 30 μg/ml gentamicin (Cytogen, 06-03100), 15 ng/ml amphotericin B (Sigma, A2942) and 100 I.U. heparin. Cells were maintained at 37°C in a humidified, 5 % CO_2_ atmosphere up to 5 passages. The cells were routinely tested for mycoplasma contamination by polymerase chain reactions.

### Plasmids and siRNA

HUVEC were transiently transfected with the plasmid DNA or siRNA using electroporation as described before ^[66]^. In brief, the Amaxa nucleofection system (Lonza) was used for electroporation according to the manufacturer’s specifications using the program U-001 with the following modifications per transfection: 1-5 μg plasmid DNA or 100-150 nM siRNA per 20 cm^2^ of nearly confluent cells were resuspended in self-made transfection buffer (4 mM KCl, 10 mM MgCl_2_, 10 mM sodium succinate, 100 mM NaH_2_PO_4_, pH 7.4 adjusted with NaOH or HCl). Cells were used for the wounding assays 18-24 h post-transfection.

The following plasmids were purchased from Addgene: LAMP1-mGFP was a gift from Esteban Dell’Angelica (# 34831) ^[67]^, EGFP-Rab7A was a gift from Qing Zhong (# 28047) ^[68]^, EGFP-Rab4A was a gift from Marci Scidmore (# 49434) ^[69]^, EGFP-Rab8a-wt was a gift from Lei Lu (# 86075) ^[70]^, pmTurquoise2-Golgi was a gift from Dorus Gadella (# 36205) ^[71]^, mApple-LAMP1-phLuorin-N-8 was a gift from Michael Davidson (# 54918), pEGFP-n1-APP was a gift from Zita Balklava & Thomas Wassmer (# 69924) ^[72]^, TfR-pHuji was a gift from David Perrais (# 61505) ^[57]^, CMV-R-GECO1.2 was a gift from Robert Campbell (# 45494) ^[73]^.

VAMP2-SEP ^[74]^, VAMP3-GFP, and VAMP8-GFP ^[75]^ were described before. GFP-2xFYVE was kindly provided by Martin Bähler (University of Münster, Germany) ^[76]^ and GFP-CD63 was procured from Jean Gruenberg (University of Geneva, Switzerland) ^[77]^.

TfR-SEP was generated by amplifying super ecliptic pHluorin (SEP) from synaptopHluorin ^[74]^, using the following primers and restriction sites and subcloning it into TfR-pHuji by replacing pHuji:

Fwd_TfR: ctcggtgatcatagttg (AgeI)
Rev_TfR: agctccctgaatagtc (XbaI)

VAMP4-SEP and VAMP7-SEP were generated by replacing VAMP2 from VAMP2-SEP with VAMP4 or VAMP7 from Syn-VAMP4-SEP and Syn-VAMP7-SEP respectively ^[78] [79]^ via restriction digestion-ligation based cloning (using BamHI and XbaI). mApple-2xFYVE was generated by exchanging EGFP from pEGFP-C2-2xFYVE with mApple from mApple-N1 using BsrgI and AgeI restriction digestion-based cloning and provided, together with GFP-Rab11b by Anna Holthenrich (University of Münster, Germany). GFP-Vti1a and GFP-DNAJC5 were generated by amplifying the corresponding sequences from a HUVEC cDNA library and cloned each into pEGFP-C1 vector (Clontech); both were provided by Johannes Nass (University of Münster, Germany). GFP-Rab5 was provided by Alexander Kühnl (University of Münster, Germany).

For protein knockdown assays, 1×10^6^ HUVEC were transfected with the corresponding siRNA (150 nM for VAMP2 and 100 nM for VAMP3) twice for 48 h and the cells were analysed 24 h after the 2nd round of transfection. The following siRNA against human VAMP2 were purchased from Dharmacon: ON-TARGETplus SMARTpool Human VAMP2 siRNA (L-012498-00-0005) and ON-TARGETplus Custom VAMP2 duplex (CTM-578530). The following siRNAs against human VAMP3 purchased from Sigma Aldrich were used: siVAMP3 #1 (NM_004781.3, 2169–2189, 5′-CACUGUAAUCACCUAAAUAAA-3′) and siVAMP3 #2 (NM_004781.3, 1462–1482, 5′-CCCAAAUAUGAAGAUAAACUA-3′). As control, non-targeting AllStars Negative Control siRNA (Qiagen Hilden, 1027281) was used.

### miRNA expression vectors

For the miRNA-mediated knockdown of VAMP2, different pre-miRNA sequences targeting human VAMP2 (GenBank accession no. NM_014232) were designed by Invitrogen’s RNAi designer (Invitrogen, 10336022). The top and bottom single strand oligos used in this study (top strand: TGCTGCAGTATTTGCGCTTGAGCTTGGTTTTGGCCACTGACTGACCAAGCTCACGCAAATACTG and bottom strand: CCTGCAGTATTTGCGTGAGCTTGGTCAGTCAGTGGCCAAAACCAAGCTCAAGCGCAAATACTGC) encode the sense and antisense strands of the pre-miRNA against VAMP2. The complementary DNA oligos were annealed to generate a double-stranded oligo and cloned into the linearized pcDNA6.2-GW/EmGFP expression vector (using the BLOCK-iT Pol II miR RNAi Expression Vector kit, Invitrogen, K493600). As a negative control, pcDNA6.2-GW EmGFP-miRneg plasmid from the kit was used. This encodes an insert that is predicted not to target any known vertebrate gene. These vectors allowed for co-cistronic expression of the EmGFP (Emerald Green Fluorescent Protein) reporter gene and synthetic miRNA under the transcriptional control of the constitutive CMV promoter. All vectors were transformed into One ShotH TOP10 Chemically Competent E. coli provided with the kit and the colonies containing spectinomycin-resistant transformants were analysed for the desired expression clones via sequencing.

For the knockdown, 1×10^6^ cells were transfected with 5 μg of EmGFP-miRNA VAMP2 or EmGFP-miRNA control plasmids and analysed after 12-18 h for knockdown efficiency using western blotting, flow cytometry, and fluorescence microscopy.

### Antibodies and reagents

The following primary antibodies were used for immunofluorescence (IF) and western blotting (WB): rabbit α-VAMP2 (1:200 IF & 1:150 WB, Proteintech, 10135-1-AP), rabbit α-VAMP3 (1:200 IF & 1:1000 WB, Proteintech, 10702-1-AP), rabbit α-VAMP4 (1: 200 IF, Synaptic Systems, 136 002), rabbit α-VAMP8 (1: 200 IF, Synaptic Systems, 104 302), mouse α-EEA1 (1:200 IF, BD Trans. Lab, 610456), mouse α-Calnexin (1:700 WB, BD Trans. Lab, 610524), rabbit α-alpha-Tubulin (1:1000 WB, Cell Signalling, 2125). For FACS based assays, rabbit α-VAMP2 Alexa Fluor 647 antibodies (EPR12790) (1:100, Abcam, ab198949) and the corresponding isotype control rabbit α-IgG (1:100, Abcam, ab199093) were used.

Immunofluorescence stainings employed the following secondary antibodies: goat α-rabbit Alexa Fluor 488 (Thermo Fisher Scientific, A11034) and goat α-mouse Alexa Fluor 568 (Molecular Probes, A-11031), whereas the following secondary antibodies were used for western blots: goat α-rabbit IRDye 800CW (LICOR, 926-32211), goat α-rabbit IRDye 680CW (LICOR, 926-32221), goat α-mouse IRDye 680CW (LICOR, 926-32220).

Lysosomal exocytosis was induced by 2 μM ionomycin (Cayman Chemical, 11932) in media. To study inhibition of lysosomal exocytosis, U18666A (Sigma-Aldrich, U3633) was used at 2 μg/ml in media for 18-24 h. Acid sphingomyelinase was blocked by desipramine hydrochloride (Sigma-Aldrich, D3900-1G) at 75μM in media for 1 hour before imaging/detection assay.

### Plasma membrane repair assays

#### Two-photon laser ablation assay

A laser ablation protocol was applied to perform membrane wounding as explained previously ^[35]^. In short, HUVEC were seeded onto collagen I-coated 8-well glass-bottom μ-slides (Ibidi, 80827) and imaged 18-24 h after transfection (50-70 % confluency). Cells were washed with prewarmed Tyrode’s buffer (140 mM NaCl_2_, 5 mM KCl, 1 mM MgCl_2_, 10 mM glucose, 10 mM HEPES, pH 7.4) ^[80]^ supplemented with either 2.5 mM Ca^2+^ or 100 μM EGTA as indicated and imaged in the same buffer containing 5 μg/ml FM4-64 (Thermo Fisher Scientific, T13320). Cells were imaged with an LSM 780 confocal microscope (Zeiss) using 63× NA 1.4 oil immersion objective (Plan-Apochromat) and a microscope stage maintained at 37°C. Laser ablation was performed with a Chameleon Vision NLO pulsed laser (Coherent) at 820 nm wavelength with the power set to 16.5 %. Using this two-photon laser, a circular region of the plasma membrane of 2.5 μm^2^ surface area (20 pixels in diameter) was irradiated for two iterations with a pixel dwell of 77 μs in “zoom bleach” mode. A time series of 2 frames were recorded before wounding to normalize for any bleaching due to imaging and a total of 100 frames was recorded to observe the wound resealing dynamics (each frame varied from 1.48-2.2 s) with 512 × 512 pixels image dimensions. Image acquisition and laser ablation were controlled with Zeiss’ Zen software.

All analyses were performed on the original, unmodified image data using Fiji ^[81]^ with the aid of custom-written scripts that are available on request. For time-lapse recordings, all fields in which cells retracted from the dish or otherwise drifted out of focus in *xy* or *z* planes during imaging were omitted from the analysis.

For FM dye influx analysis, a macro utilizing the “Plot Z-profile” function of Fiji was developed as detailed in ^[36]^. A maximum intensity projection of the FM4-64 channel was created for the acquired time series and the first frame prior to wounding was subtracted from all other frames within the time series. The analysis was proceeded by outlining the wounded cell and an uninjured cell for background correction (to compensate for any endocytic uptake of FM4-64 during acquisition). The kinetics of plasma membrane repair was determined by FM4–64 fluorescence intensity change (absolute units, with the background correction) over the entire wounded cell as a function of time. Membrane repair was determined by the entry of FM dye into the cell interior, wherein a plateau in FM dye increase indicated successful repair, and failure to repair was indicated by a linear increase in the FM dye influx.

#### Glass bead injury assay

Mechanical injury using glass beads was performed as outlined ^[37]^ with a few modifications. A total of 1.6 × 10^5^ siRNA transfected or untransfected cells were cultured on collagen-coated 2-well glass-bottom μ-slides (Ibidi, 80287) until 95-100 % confluency (20-24 hours).

Untransfected cells were pretreated with the drugs, U18666A (2 μg/ml for 18-22 h) or desipramine (DPA, 75 μM for 1 hour) or corresponding vehicle controls before glass bead injury. Cells were transferred to prewarmed phosphate buffered saline (PBS+/+) supplemented with Dextran-Alexa Fluor (AF) 488 (1 mg/ml, Thermo Fisher Scientific, D22910) and injured by rolling 300 mg of acid-washed glass beads (425–600 μm, Sigma-Aldrich, G8772-100G) over the cells for 1 min. Cells were then allowed to reseal in PBS+/+ at 37°C for 5 min. The cells were further incubated with PBS+/+ supplemented with Dextran-Alexa Fluor 546 (1 mg/ml, Thermo Fisher Scientific, D22911) for 1 min. Following this, the cells were fixed with 4 % paraformaldehyde (PFA) and nuclei were counterstained with Hoechst 33342 (40 μg/ml, Thermo Fisher Scientific, 62249) for 30 min. The slides were then imaged using an LSM800 confocal microscope (Zeiss) equipped with a 20× objective. A series of at least 20 different fields were recorded for each sample. The number of AF488-positive cells (injured and repaired) and AF546-positive cells (injured and non-repaired) were counted post-thresholding using the “Cell Counter” plugin in Fiji. The efficiency of membrane repair was represented as a percentage of the repaired cells to the total number of beads-injured cells. It is important to note that the number of bead injured cells varied across samples due to variability in wounding efficiency ^[49] [50]^. Therefore it is essential to compare the effective resealing ratios with the corresponding controls.

The assay was validated for defects in membrane repair in the absence of Ca^2+^ by rolling the beads in the presence of PBS−/−, but quantification of repair efficiency was difficult due to severe cell loss via detachment of HUVEC from the dish (see also Figure S1).

#### Mechanical scrape assay

Mechanical injury by scraping was performed with adaptations from ^[48] [82]^. As with the glass bead injury, 5-6 × 10^5^ siRNA transfected or untransfected cells were cultured on collagen-coated 6-well plates (Ibidi, 80287) until 90-100 % confluency (20-24 hours). Before scraping, untransfected cells were pretreated with the drugs as specified above. Cells were washed with prewarmed PBS+/+ or PBS−/− and then kept in the corresponding PBS supplemented with Dextran-Alexa Fluor 488 (200 μg/ml, Thermo Fisher Scientific, D22910). Using a cell scraper (Sarstedt, 83.3951), cells were injured rapidly over the entire well in a gentle and consistent pattern to ensure that the entire well area was covered once and to maintain reproducibility in wounding. The injured cell suspension was gently pipetted into a pre-warmed Eppendorf tube and allowed to reseal at 37°C for 5 min. Following this, the repair reaction was immediately stopped in ice and the samples were incubated with ice-cold propidium iodide (PI) (50 μg/ml, Sigma Aldrich, P4864-10ML) for 4 min in ice. The cell suspension was then collected at 200 *g*, 4 min, 4°C, resuspended in Cell Wash buffer (BD Biosciences, 349524), and analysed by flow cytometry (Guava easyCyte 11 Flow Cytometer, Merck). A total of 5000-10000 cells were measured for each condition, analyzing the fluorescence with the 488 nm laser line and emission at 525/30 nm for AF488 and 695/50 nm for PI. The fragmented cell debris and aggregates generated due to the assay were segregated with a strict gating protocol using a control sample of uninjured cells stained with propidium iodide. This helped to assure that only the population with whole cells was selected (along with the forward and side scatter of uninjured cells). The cells treated with the inhibitors suffered general cell loss due to detachments during the drug treatments and subsequent scraping and thus, measurements were pooled from multiple technical replicates for each biological experiment. The AF488-positive cells (injured) indicated the efficiency of injury and PI-positive cells among the AF488-positive cells outside the gated region (injured and non-repaired) cells indicated the non-resealed cells. Data analysis was performed using InCyte software (Guava). The number of repaired cells was determined from the fraction of PI negative cells within the gated region (dashed lines) and within the total injured cells (based on AF488 fluorescence) per sample. The efficiency of membrane repair was represented as a percentage of the repaired cells to the total number of scrape-injured cells. Again, as outlined in the glass bead assay, it is important to compare the effective resealing ratios with the corresponding controls to ensure no interpreting bias.

#### Staining of intracellular compartments

HUVEC were stained for different intracellular compartments using the following markers:

Endoplasmic reticulum (ER): ER-Tracker Red (BODIPY™ TR Glibenclamide) (Thermo Fischer, E34250) was added to cells at 1 μM for 30 min at 37°C in Na-Tyrode’s buffer. The solution was then washed off before imaging.

Lysosomes: LysoTracker Green DND-26 (Thermo Fischer, L7526) was added at 150 nM for 30 min at 37°C in Na-Tyrode’s buffer to cells. The solution was replaced with fresh buffer before imaging.

Mitochondria: MitoTracker Green FM (Thermo Fischer, M7514) was added to cells at 0.1 μM for 30 min at 37°C in Na-Tyrode’s buffer. The solution was then washed off before imaging.

#### Lysosomal exocytosis assay

HUVEC were transfected with the lysosomal membrane protein fusion construct, pHluorin-LAMP1-mApple, and cultured on collagen-coated 8-well glass-bottom μ-slides for 20-24 hours. For the ionomycin-based exocytosis assay, cells were washed and imaged in endothelial mixed media supplemented with 20 mM HEPES. LAMP1 fluorescent protein-expressing cells were imaged live at the basal membrane plane (similar to ^[83]^) for 5 frames to obtain the basal expression and secretion of LAMP1 at the membrane surface and then ionomycin (2 μM final concentration) was added to induce Ca^2+^ influx and thereby lysosomal exocytosis. A time series for 2 min was recorded to ensure sufficient time for ionomycin mediated Ca^2+^ influx which results in lysosomal exocytosis ^[42]^. The latter was detected as the appearance of bright punctae in the pHluorin channel over time and each punctae was corroborated to be lysosomal exocytosis events with the LAMP1-mApple expression at the same spatial position. For these experiments, a focus shift was observed in certain images during DMSO or ionomycin addition (due to the opening of the heating chamber and the solution influx) and these frames were omitted for analysis. The first frame with the correct plane of LAMP1 fluorescence signal after DMSO or ionomycin addition and the consecutive frames were used for punctae detection across all samples.

For the laser wounding assay, pHluorin-LAMP1-mApple expressing cells were imaged in Tyrode’s buffer with FM4-64 (as in the membrane repair assay) and cells were wounded. The wounding process triggers Ca^2+^ influx that induces lysosomal exocytosis, which again can be observed as bright punctae at the membrane similar to the ionomycin assay. For the inhibition of lysosomal exocytosis, the cells were pretreated with U18666A and assessed for efficiency of treatment by bright field imaging of the cells which show an accumulation of enlarged lysosomes near the perinuclear region of the cells. Following this, the above assays were performed in the presence of U18666A. Effective lysosomal exocytosis was quantified by creating probability maps using the pixel classifier software iLastik ^[84]^ following a thresholding step in Fiji (*xy*) to obtain distinct LAMP1-pHluorin signals. The pHluorin punctae following ionomycin treatment or wounding were counted over time over the entire cell, normalized to the basal LAMP1-pHluorin expression, and represented as a percentage.

#### Acid sphingomyelinase (ASM) activity assay

ASM activity was measured using the Amplex Red sphingomyelinase Assay Kit (Thermo Fisher Scientific, A12220). In brief, HUVEC were grown on 10 cm Corning dishes for 20-24 h to confluence and treated with the ASM inhibitor, DPA, or water control for 1 hour at 37°C. After treatment, the cells were washed twice with ice-cold PBS and placed on ice. For the lysate preparation, the cells were harvested by scraping in acidic buffer (50 mM sodium acetate, 150 mM sodium chloride, pH 5.0) supplemented with protease inhibitors (Roche, 11873580001) to ensure maximum retrieval of ASM enzyme and then subjected to freeze-thaw cycles in liquid nitrogen thrice. The lysed cells were sonicated, and the cellular debris was removed by centrifugation at 15,000 *g* for 5 min at 4°C. The protein concentration in the supernatant was determined using Pierce 660 nm protein assay (Thermo Fisher Scientific, 22660) and the amounts of protein (~0.5 μg) in all samples were normalized to equal amounts. The ASM activity assay was then performed according to the manufacturer’s protocol on a 96-well black bottomed plate (Greiner, 655076) and the measurements were recorded using a fluorescence plate reader (Clariostar, BMG LabTech) maintained at 37°C over time. The purified sphingomyelinase enzyme supplied by the manufacturer served as a positive control to ensure the specificity of the assay. Similarly, the robustness of the assay was validated by the H_2_O_2_ only sample which served as a negative control. The fluorescence was measured over time using excitation at 545 ± 20 nm and fluorescence detection at 600 ± 40 nm. The readings were taken till the measurements reached a plateau and this time point was used for the calculation of the effective enzyme activity. The values were corrected for the background signal that was determined in the samples treated in the same way as described above but not containing cell lysate. The effective ASM activity was represented as a percentage normalized to the enzyme activity in the control untreated cells.

#### Immunocytochemistry

Endogenous localization of the various VAMP proteins (VAMP 2/3/4/8) was determined using immunofluorescence. Shortly, HUVEC were grown on collagen-coated 12 mm diameter coverslips. After reaching 70-80 % confluency, cells were fixed in 4% PFA for 10 min at room temperature and permeabilized using 0.1 % Triton-X-100 for 2 min. Cells were then blocked in 3% BSA in PBS for 30 min and subsequently stained with the respective VAMP or early endosomal antibodies for 1 hour. Following PBS washes, the cells were stained with the corresponding fluorescent dye labelled secondary antibodies for 40 minutes and then mounted in mowiol. Imaging employed an LSM 800 laser scanning microscope (Zeiss) equipped with a 63x (NA 1.4) oil immersion objective (Plan-Apochromat). Z-stack projections of at least 10 cells were recorded per condition.

#### Western blotting

HUVEC grown to confluence on 10 cm collagen-coated dishes were washed with ice-cold PBS and then scraped on ice into lysis buffer (50 mM Tris pH 7.4, 150 mM NaCl, 1 mM EDTA, 1 % NP-40, 0.5 % Na deoxycholate) containing protease inhibitors. The lysates were sonicated for 1 min at 4°C and the cellular debris was removed by centrifugation at 10000 *g* for 4 min at 4°C. The protein concentration in the cleared lysate was determined by Pierce 660 nm protein assay. After normalization of protein content, samples were boiled in Laemmli buffer at 95°C for 10 min. Equal amounts of lysates were subjected to SDS-PAGE and western blotting according to standard protocols ^[85] [86]^. The proteins were resolved by SDS-PAGE employing 12 % or 15 % PAA gels depending on their molecular mass with electrophoresis at 80 V for 30 min and then at 110 V till the protein ladder was well separated. The proteins were then transferred to 0.2 μm nitrocellulose membranes (Millipore, 10600001) using a wet tank system at 115 V for 1 h at 4°C in Tris-Glycine buffer (25 mM Tris, 190 mM glycine, 20 % (v/v) methanol). The membranes were then further blocked in 5 % milk in 1x Tris-buffered saline-Tween (150 mM NaCl, 50 mM Tris-HCl, pH 7.6) supplemented with 0.1 % Tween-20) (TBST), for 1 hour and probed with primary antibodies overnight at 4°C. The next day following TBST washes, secondary antibodies conjugated with infrared dyes were used (1 h staining), and the proteins were detected using the Odyssey infrared imaging system (Li-COR). The efficiency of protein knockdown was quantified in Fiji using the “Gel Analyzer” tool and represented as a percentage of the ratio of the respective protein signal intensity against the loading control.

#### miRNA knockdown detection by flow cytometry

EmGFP-miRNA control or EmGFP-miRNA VAMP2 transfected HUVEC were seeded onto collagen-coated 6-well dishes. After 12-18 h, cells were harvested by trypsinization, then washed with PBS, fixed, and permeabilized with BD Cytofix/Cytoperm (BD Biosciences, 554714) for 20 min under light exclusion conditions. Following centrifugation at 200 *g* and 4°C, the cells were blocked with BD Perm/Wash buffer (BD Biosciences, 554714) for 30 minutes under light exclusivity and spun down again. The resulting pellet was incubated with anti-VAMP2 primary antibodies conjugated to Alexa Fluor 647 for 1 hour in darkness. The cells were washed thrice with Cell Wash and then resuspended in Cell Wash and analysed by flow cytometry. The samples were gated according to the cell size and population by their forward and side scatter properties. Following this, 10000 cells positive for GFP fluorescence were counted per sample, indicating successful miRNA transfection. These cells were gated and analysed for VAMP2-AF647 staining. Data analysis was performed using InCyte software (Guava). The median fluorescence intensity of VAMP2 staining was determined based on the total number of GFP-positive cells. The efficiency of knockdown was depicted as a ratio of the VAMP2 median fluorescence intensity in miRNA VAMP2 samples normalized to the miRNA control.

#### Correlative Light and Electron Microscopy (CLEM)

HUVEC were seeded on glass-bottomed dishes (Ibidi, 81218-200), specifically coated with a carbon pattern to re-locate the cell of interest by electron microscopy (Finder grid mask, Leica, see also Figure S13), and cultivated to reach about 70-80 % confluency. Prior to the experiment, cells were labeled with BSA-Au (CMC, Utrecht, BSAG 10nm) for 90 minutes in medium. After a brief wash, Tyrode’s buffer supplemented with FM4-64 membrane dye (used for laser ablation assay) and Ca^2+^ or EGTA as indicated, was added to the cells. Subsequently, the samples were immediately processed at the LSM 780 microscope for the laser wounding experiment as detailed above. One cell per dish next to an identifiable pattern was selected and documented with respect to the carbon grid for later relocation. Care was taken to ensure that the cell was suitable for the laser ablation assay and had landmarks nearby for relocation. Images were acquired at 20x and 63x magnification for this purpose next to the carbon coated pattern. Following wounding, the cell was imaged for 40 seconds, and immediately thereafter the sample was pre-fixed by adding 1 volume of 4 % PFA in 0.1 M PHEM buffer resulting in 2 % paraformaldehyde prefixation conditions. During this time, the membrane labelling of the wounded cell over the entire cell volume was imaged by acquiring a *z*-stack with thin intervals to obtain detailed information about the wound site and cell size. The solution was changed after 10 minutes to 2 % glutaraldehyde + 2 % PFA in 0.1 M cacodylate buffer for the final fixation. Samples were flat embedded in epon starting with post-fixation in 1 % osmium tetroxide including 1.5 % potassium cyanoferrate. Dehydration was done in ethanol with a 0.5 % uranyl acetate *en*-bloc staining during the 70 % ethanol step. Stepwise infiltration in mixtures of ethanol and epon followed until pure epon was reached. For flat embedding, the area of interest was finally covered with a gelatin capsule filled with epon. The blocks were polymerized for 3 days at 60°C. The glass bottom was detached in liquid nitrogen so that the carbon marks remained etched on the block surface and could be used as a landmark to relocate the cell of interest. The surface of the block was reduced to a square of a maximum of 500 μm x 500 μm, retaining the cell of interest in the center. Sectioning with a diamond knife ensued directly and all sections were taken (60 nm ultrathin and 250 nm thick sections) from the basolateral side to the apical side of the cell and collected on formvar coated grids. The grids were stained with lead and analysed and imaged at a transmission electron microscope (Tecnai12-biotwin, Thermo Fisher Scientific, Netherlands) with careful correlation to the fluorescence signal recorded after wounding.

Images from 250 nm thick sections labelled with 15 nm gold fiducials (BBI Solutions, UK) were obtained by using a JEOL JEM-2100Plus 200 kV transmission electron microscope (JEOL, Japan) equipped with a XAROSA CMOS 20 Megapixel camera (EMSIS GmbH, Germany). The pixel size was 1.17 nm with a nominal magnification of 8000x or 0.79 nm at 12000x respectively. One or two perpendicular tilt series per tomogram with an angular range of ±60° with 1° increments were acquired and a dual-axis tomographic reconstruction was computed using the back projection algorithm in IMOD ^[87]^. For the chosen regions of interest, cellular objects, mainly endosomes and plasma membrane, were selected and contoured throughout the 3D volume, resulting in a model with 3dmod. Finally, the closed contours were meshed, and the *z*-scale was stretched with 1.4 to correct for resin shrinkage. The final movies were generated using Fiji.

#### Detection of early endosome exocytosis by STED

Exocytosis of early endosomes in wounded cells was detected using a pHluorin-nanobody based assay as described before ^[58]^. In brief, HUVEC transfected with TfR-SEP or VAMP2-SEP were grown on collagen-coated gridded 35mm μ-dishes (Ibidi, 81148) for 24 hours. Cells were washed and blocked with an equal mixture of unconjugated FluoTag-X2 anti-GFP nanobodies (Nanotag Biotechnologies, C1000, clones 1H1 and 1B2) at 1:20 dilution in Na-Tyrode’s buffer with Ca^2+^ for 2 minutes at 37°C. Following this, the cells were washed and subjected to laser ablation at the LSM 780 microscope. A transfected cell next to a distinct pattern on the gridded dish was selected and the location was noted. Just before wounding, FluoTag-X4 anti-GFP nanobody labelled with Abberior Star 580 (Nanotag, N0304-Ab580) at 1:50 dilution was added to the dish. The laser ablation protocol was performed as described above (omitting FM-464 dye) and immediately (1.5 s) after wounding, the dish was fixed at room temperature with 2 % PFA for 3 minutes followed by 4 % PFA for 10 minutes. After a few washes with PBS, the wounded cell, relocated using the gridded dish pattern, was imaged using confocal and STED microscopy. STED images were acquired with a Leica TCS SP8 STED 3x microscope equipped with a 93×1.3 NA HC PL APO CS2 glycerol immersion objective. Excitation was performed by a pulsed (80 MHz) tunable white light laser and emission was detected by hybrid detectors. First, pHluorin and Abberior Star 580 were sequentially excited in confocal mode at 488 and 561 nm, respectively. Then, Abberior Star580 was imaged in time-gated STED mode using 561 nm excitation. The excitation spot was superimposed by a *xy* depletion doughnut generated by a pulsed 775 nm laser (1.5 W output). The intensity of the depletion beam was between 50-100 % depending on signal intensity and the pixel size was set to 40 nm x 40 nm. The resulting images were filtered with a Gaussian blur of 1.0-pixel size and displayed with equal linear contrast adjustment across samples.

#### Quantification of endosomal punctae on wounding

HUVEC were transfected with the various early and late endosomal markers or pulsed with Transferrin-Alexa Fluor 488 (Thermo Fisher Scientific, T13342) for 5 min at 37°C, and laser wounding was performed as stated above. A time series of 2 frames were recorded before wounding to normalize for any bleaching and a total of 100 frames was recorded to observe the wound resealing dynamics (each frame varied from 1.48-2.3 s depending on the fluorescence channels). The images were subsequently analysed in Fiji using a macro developed by Sophia Koerdt (University of Münster, Germany) to quantify the change in the localization of endosomal markers. The images were background-subtracted using a rolling ball radius, filtered, and thresholded to binarized images. To discern the punctate localization of endosomes, a size threshold below an empirically determined size (over various punctae across cells) was used. The same threshold was then applied to the recordings of all samples analysed within each experiment.

Following this, an ROI (region of interest) - based analysis approach was applied. Specifically, a set of concentric circular ROI of increasing radii (20 μm each in diameter) was defined with respect to the circular wound ROI (2 μm^2^). This separated the wounded cell into different zones in distance from the wound site, e.g., circle 1 being closest to the wound site and circle 5 being the farthest from the wound ROI. The number of punctae in each of these circular ROIs was measured over time for the different endosomal markers. Further, the number of puncta identified at each time point was normalized to the initial number of puncta, and calculated as the average of the first two time points. Additionally, to identify the punctae at the cell edge of the HUVEC membrane, a circular ROI (10 μm in diameter) was defined from the boundary of the cell, termed circle 6. These punctae measurements of the markers were represented as a percentage of the disappearance of endosomal punctae over time. A similar analysis was applied to a non-wounded control where cells were ablated with the two-photon laser at 0.2 % laser power. The number of punctae quantified in these cells was used to set the baseline for the dynamics of the endosomal markers through *z* and time.

For the quantification of the correlation of disappearing 2xFYVE punctae with the Ca^2+^ indicator R-GECO 1.2, similar analyses were employed, except the intensity measurements of the Ca^2+^ indicator were calculated on raw non-binarized images for each circular ROI from the wound site over time. Additionally, thresholding of the 2xFYVE punctae was applied by first creating probability maps using the pixel classifier software iLastik followed by thresholding the cells in Fiji (*xy*) prior to the count analysis. The dynamics of the Ca^2+^ wave, represented as an increase in signal from the baseline expression, was correlated to the localization of 2xFYVE punctae in each circle ROI over time after wounding.

#### Colocalization analysis

Colocalization of VAMPs with early endosomal markers was determined as follows. Cells were pulsed with Transferrin-Alexa Fluor 647 (Thermo Fisher Scientific, T23366) in media at 37°C for 5 min before fixation (to label transferrin cargo localized to early endosomes). Transferrin uptake was then stopped on ice followed by ice-cold PBS washes. The cells were then fixed in 4 % PFA and the rest of the immunofluorescence protocol was carried out as outlined above. An anti-EEA1 antibody was used as an additional early endosomal marker along with the anti-VAMP antibodies during primary antibody staining. During imaging, *Z*-stack projections of at least 10 cells were recorded for each VAMP protein per experiment. The degree of colocalization between the different VAMPs and each of the early endosomal markers was then analysed using the JaCoP plugin ^[88]^ and pixel by pixel covariance was measured in Fiji. For this, the cell of interest was outlined and thresholded before Pearson’s correlation coefficient, r, was calculated. The *r*-value was then plotted for the different VAMPs with reference to each early endosomal marker (transferrin and EEA1).

#### Detection of early endosome exocytosis events

HUVEC were cotransfected with the different VAMP proteins (VAMP2-SEP, VAMP3-GFP, VAMP4-SEP, or VAMP8-GFP) or GFP-2xFYVE along with the endosomal membrane marker, TfR-pHuji, and cultured on collagen-coated 8-well glass-bottom μ-slides for 20-24 hours. Membrane wounding was performed on the different VAMP/2xFYVE and TfR cotransfected cells as detailed in the laser ablation assay. A time series of 2 frames were recorded prior to wounding to normalize for any bleaching and a total of 40-100 frames was recorded to observe the wound resealing dynamics (each frame varied from 1.48-2.3 s). Exocytosis events were detected as bright SEP/pHuji accumulations/clusters distinct from the background over a time series of 30 frames. As before, the images were subsequently analysed using iLastik and Fiji. iLastik was used to create probability maps by using supervised machine learning approaches across all images to generate probability maps to differentiate signal and background and ensure accurate detection of the exocytotic clusters. Thresholding of the resulting probability maps was performed in batch for all images using a Fiji macro. To capture the exocytotic clusters that form at the sites of 2xFYVE disappearance in the same frame and the next frame, a walking average for 2 frames was performed to extract the maximal 2xFYVE-associated exocytosis events. Next, an ROI-based analysis approach, as described above for the endosomal punctae count, was applied to the thresholded images. In addition to the punctae count in the concentric circular ROIs, an object based colocalization analysis was performed on the thresholded and averaged frames to get the number of colocalized punctae positive for the different VAMPs/2xFYVE and the TfR-pHuji clusters. Following the measurements in Fiji, the punctae count in each concentric ROI was normalized per μm^2^ of ROI area in the cell to account for the variability in the area of the ROIs from the wound site. Furthermore, the number of punctae per μm^2^ identified at each time point was normalized to the initial number of punctae prior to wounding. It is to be noted that the null baseline values were set to 1 to ensure uniform normalization across samples, for the further quantification of exocytotic events. The increase in early endosome or VAMP exocytotic events was plotted over time after wounding.

#### Statistical analysis

All numerical values are given as means and error bars are standard deviations (SD) if not stated otherwise. Unless mentioned else in the figure legends, *N* represents the number of independent experiments, and *n* denotes individual cells obtained from *N* biological samples. In each assay, a minimum of three biological replicates was performed and the specific number of replicates per independent experiment is noted in the corresponding figure legends. Data were analysed and plotted using GraphPad Prism 8.4.3 (GraphPad Software). All experimental datasets were checked for their normality of distributions using Shapiro-Wilk test. Accordingly, parametric and nonparametric tests were chosen as appropriate for the data and are reported in the figure legends. Specific statistical details for each experiment can be found in the corresponding figure legends.

Data were considered statistically significant at the following *p*-values: <0.05 (*), <0.01 (**), <0.001 (***), <0.0001 (****). *ns* denotes not significant.

#### Data representation

All digital images, including fluorescence and electron micrographs, immunoblots, and time-lapse recordings, were linearly contrast stretched in Fiji to display relevant features. All frames within a movie, and all images and movies intended to be compared, were processed identically to ensure there was no bias. Data were plotted using GraphPad Prism 8.4.3. Figures were assembled for publication using Fiji and Inkscape (San Jose, CA, USA). Animations were created with BioRender.com.

## Data availability

All data are available in the main text or the supplementary information. Additional data or codes related to this paper may be requested from the authors.

## Acknowledgments

We thank Sophia Koerdt (University of Münster) for developing the macro for initial image analysis, Silvio Rizzoli (University of Göttingen) for advice on the nanobody-based exocytosis assay and Tomas Kirchhausen (Harvard Medical School) for comments on the manuscript. We are indebted to Michelina Kierzek (University of Münster) for critical reading of the manuscript and to Anna Holthenrich and Anna Matos (University of Münster) for fruitful discussions. We also thank members of the Gerke lab and the Institute of Medical Biochemistry for reagents, support, and discussions. NR is a member of CiM-IMPRS, the joint graduate school of the Cells-in-Motion Interfaculty Centre, University of Münster, Germany and the International Max Planck Research School - Molecular Biomedicine, Münster, Germany. This work was supported by grants from the German Research Foundation (CRC1009, project A06, and GE514/6-3 to V.G, and CRC1944, Z project, to O.E.P.)

## Author information

V.G. supervised the project. N.R., D.Z. and V.G. conceived the study. N.R. performed the experiments. L.G., K.M., D.Z., R.F. and O.E.P contributed to the electron microscopy and tomography experiments. M.K. and J.K. contributed to STED imaging experiments. T.Z. wrote the codes for image analysis. N.R. and V.G wrote the original manuscript with contributions from D.Z., M.K and T.Z. All the authors discussed and commented on the manuscript.

## Corresponding author

Correspondence to Volker Gerke.

## Ethics declarations

### Competing interests

The authors declare that they have no competing interests.

## Notes

### Competing Interest Statement

The authors have declared no competing interest.

