## Supplementary file for "Early endosomes act as local exocytosis hubs to repair endothelial membrane damage"

**This PDF file includes:**

Figs. S1 to S22
Captions for Videos S1 to S10

**Other Supplementary Materials for this manuscript include the following:**
Videos S1 to S10

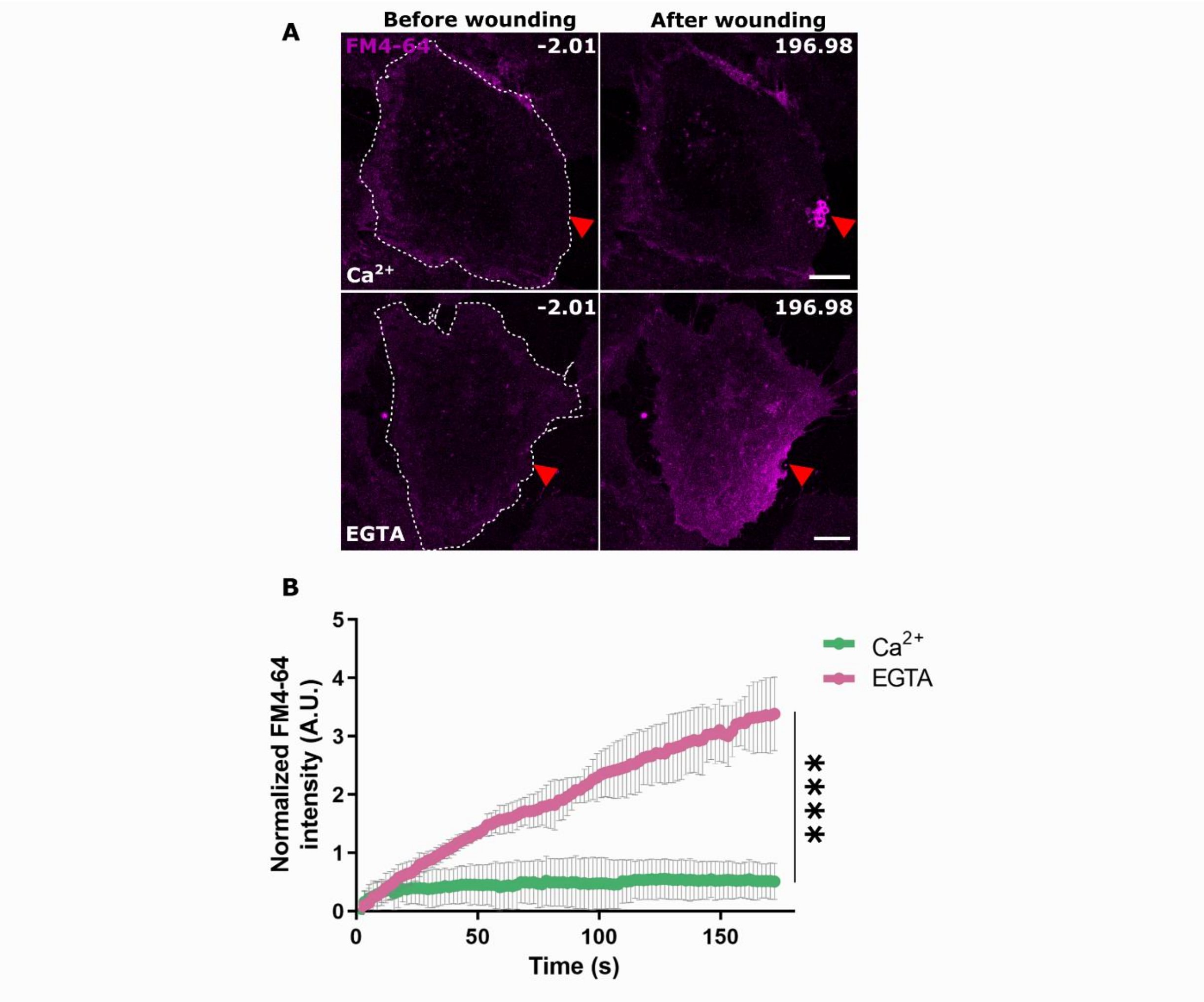


**Figure S1. Kinetics of plasma membrane resealing in HUVEC wounded by two-photon laser ablation. (A)** HUVEC were incubated in buffer containing $Ca^{2+}$ or the $Ca^{2+}$ chelator EGTA and supplemented with the membrane-impermeable fluorescent dye FM4-64 (magenta). Cells were locally wounded at the lateral edge of the plasma membrane employing a two-photon laser at 820 nm and a wound ROI of 2 $\mu m^2$. Time-lapse imaging was performed after wounding to track the kinetics of membrane resealing. Representative images of a cell pre and post wounding in the presence or absence of $Ca^{2+}$ are shown here. Red triangle, wound site. White dashes indicate the outline of wounded cells. Scale bars, 10 µm. (**B**) Graph showing the efficiency of membrane resealing as measured by the influx of the FM4-64 dye. Mean $\pm$ SD of the increase in fluorescence intensity is shown, normalized to the intensity before wounding of the same cell and a neighbouring unwounded cell. 24 cells per condition pooled from three independent experiments, $P < 0.0001$ with two-tailed Mann-Whitney $U$ test performed.

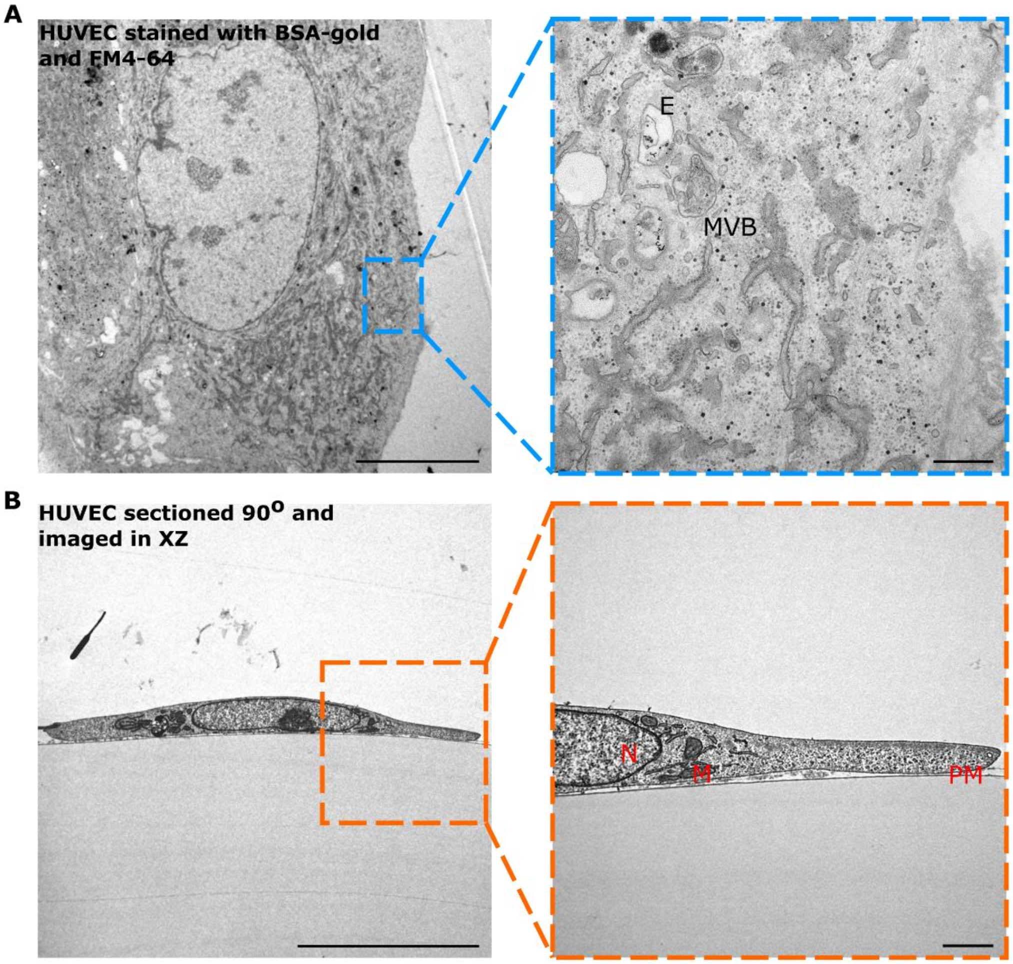


**Figure S2. HUVEC are flat cells ideal for membrane repair studies following lateral membrane laser ablation. (A)** HUVEC were labelled with BSA-gold for 90 minutes to mark all endocytic compartments, washed, incubated in media supplemented with FM4-64, fixed, and processed for TEM as described in Methods. Representative TEM image of an ultrathin 60 nm section showing that the morphology of HUVEC is unaffected by FM4-64 labelling. Higher magnification of the dashed blue area shows the efficient uptake of BSA-gold in various endosomal compartments, which are of regular size. Endosomes indicated with E, multivesicular bodies indicated with MVB. Scale bars, 10 µm; for zoom, 500 nm. (**B**) HUVEC, originally flat embedded, were re-embedded in 90° to the original orientation to visualize the cell in the XZ expansion. TEM image shows the thickness of the endothelial cell. Zoom-in of the dashed orange box reveals the thickness of the lateral plasma membrane stretch to be less than 500 nm, which was within the diffraction limit of the microscope used for laser injury. Nucleus marked with N, mitochondria marked with M, plasma membrane marked with PM. Scale bars, 10 µm; for zoom, 1 µm.

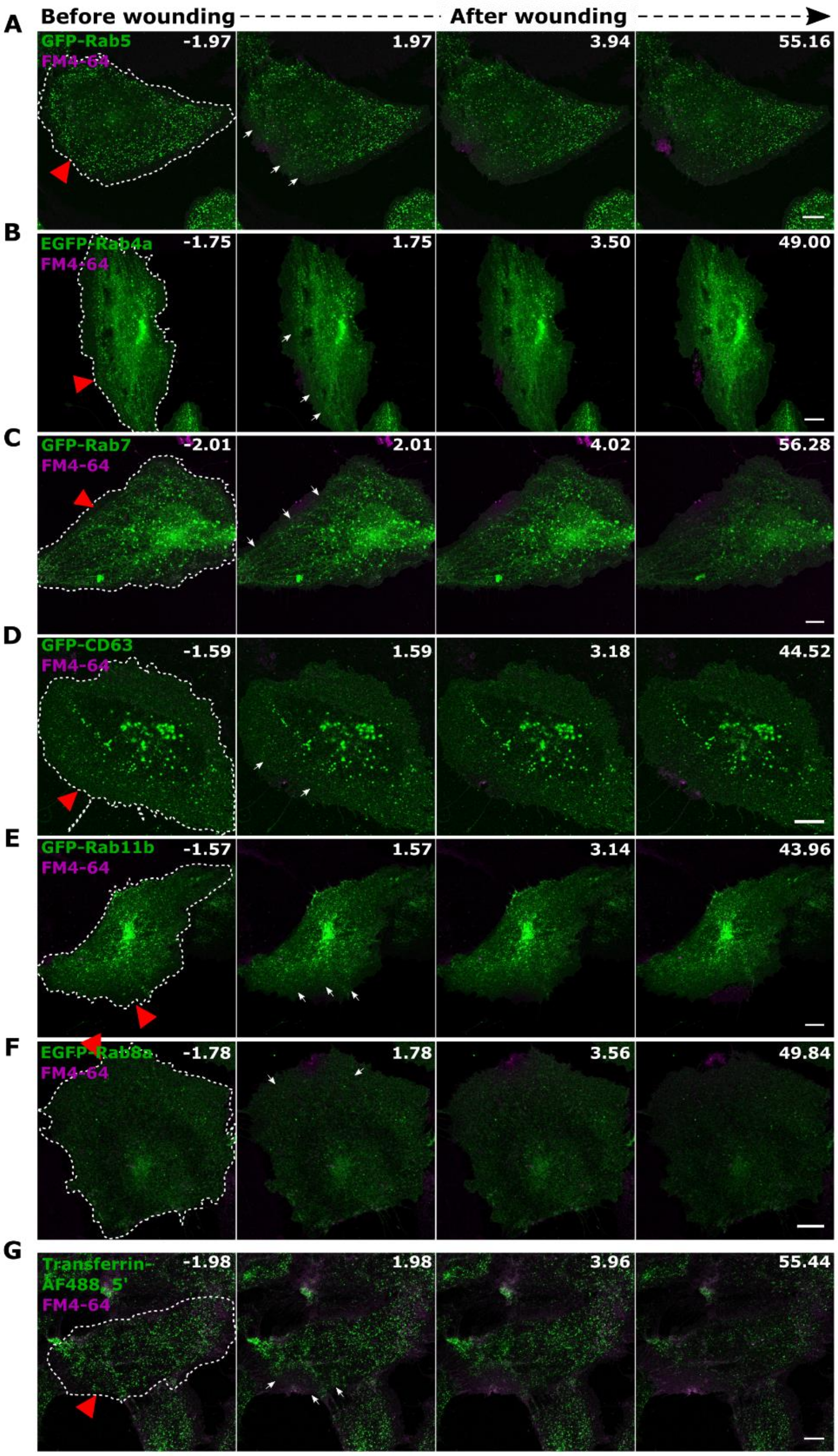


**Figure S3. Markers of endosomal compartments disappear near the wound site following PM injury in HUVEC. (A - F)** HUVEC were transfected with various endosomal compartment markers (green in all panels), GFP- Rab5 (A), EGFP-Rab4a (B), GFP-Rab7 (C), GFP-CD63 (D), GFP-Rab11b (E) or EGFP-Rab8a (F), and subjected to laser injury in the presence of FM4-64 (magenta). Representative sequential images before and after wounding are shown (t = 0 *s*

represents the time of wounding). All markers showed varying amounts of signal disappearance near the wound site immediately after wounding. A particularly pronounced disappearance near the wound site is observed for early endosomal markers (GFP-Rab5 and EGFP-Rab4a; panels A and B), with minimal disappearance observed for GFP-Rab8 or GFP-CD63 (panels D and F). Red triangle, wound site, and white dashes, wounded cells. White arrows indicate sites of endosomal disappearance after wounding. Localization of FM4-64 dye to the wound site indicated successful membrane resealing in all examples. (**G**) HUVEC were labelled with fluorescent transferrin cargo for 5 min to populate early endosomes, washed, and then wounded immediately. A marked disappearance of early endosomes positive for transferrin was observed near the wound site. All markers were Scale bars, 10 μm.

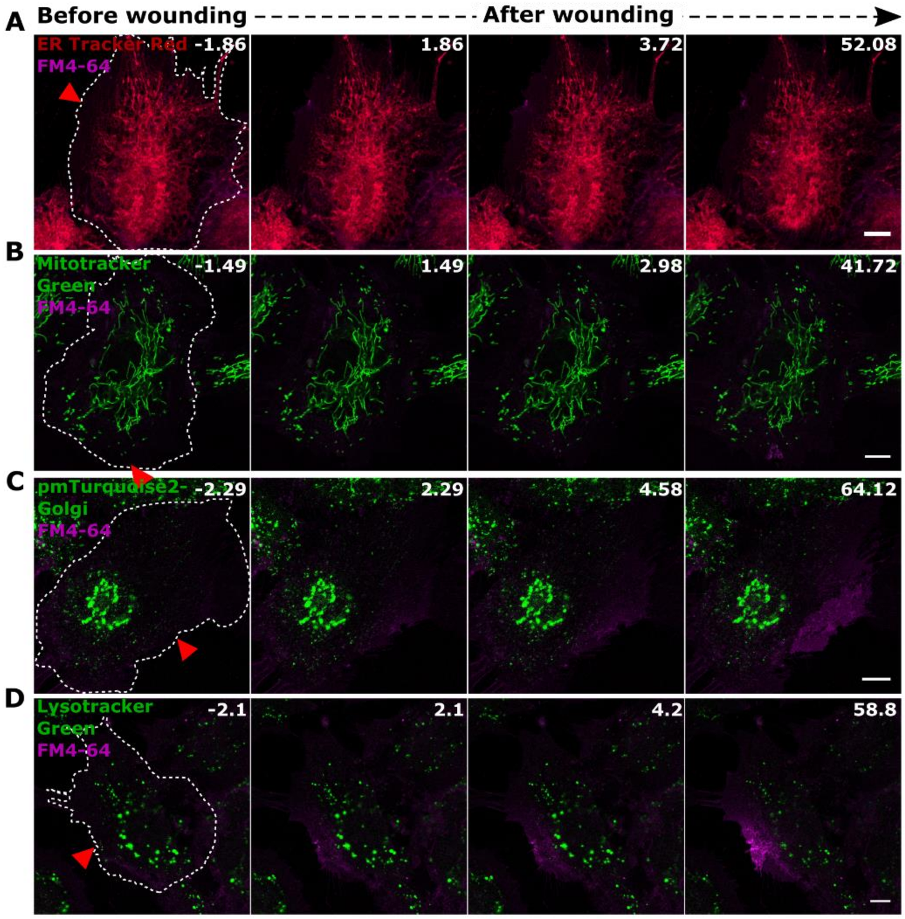


**Figure S4. Various intracellular organelles appear unaffected by PM wounding in HUVEC. (A - D)** HUVEC were stained for organelle markers – endoplasmic reticulum (ER) (shown in red) (A), mitochondria (B), and lysosomes (D) - or transfected with a Golgi marker (C) and wounded by laser ablation in the presence of FM4-64 (shown in magenta and all other markers in green unless indicated otherwise). Time-lapse images of cells before and after wounding are shown. No appreciable change in the localization of the various organelle markers was observed after wounding. Also note that the Lysotracker staining, which labels the perinuclear pool of lysosomes, showed no association with the wound site. Red triangle, wound site, and white dashes, wounded cells. Scale bars, 10 µm.

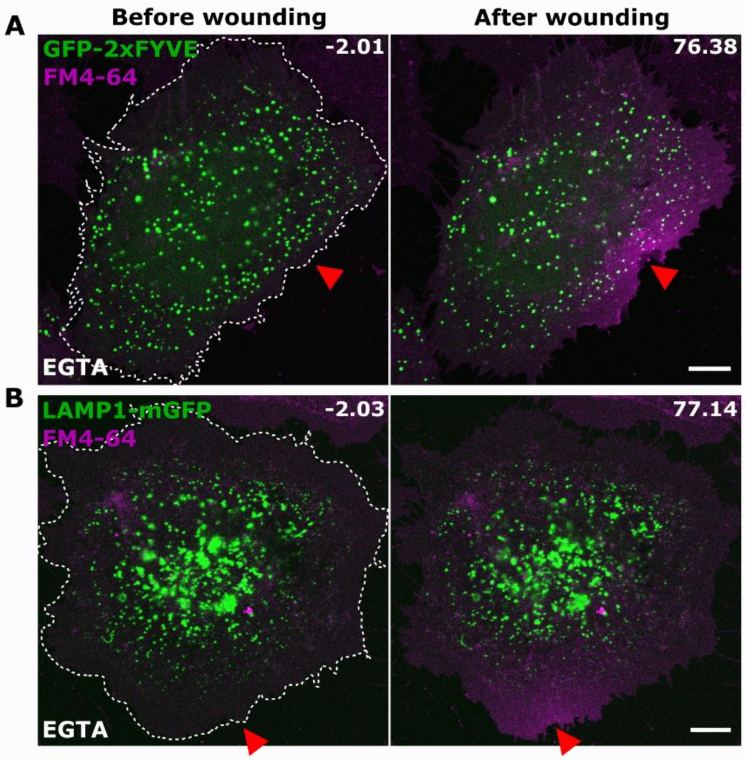


**Figure S5. Disappearance of early and late endosomal markers after membrane injury is a $Ca^{2+}$- dependent response. (A - B)** HUVEC ectopically expressing the early endosomal marker GFP-2xFYVE (A) or the late endosomal/lysosomal (LEL) marker LAMP1-mGFP (B) were subjected to laser injury in medium containing FM4-64 (magenta; everything else displayed in green) and EGTA. Representative images of the same cell before and after wounding are shown. No discernible disappearance of the endosomal markers can be observed upon wounding in the absence of external $Ca^{2+}$. Impaired membrane resealing as revealed by the increase in FM4-64 intensity indicates the inhibition of wound repair in the presence of EGTA. Red triangle, wound site, and white dashes, injured cells. Scale bars, 10 μm.

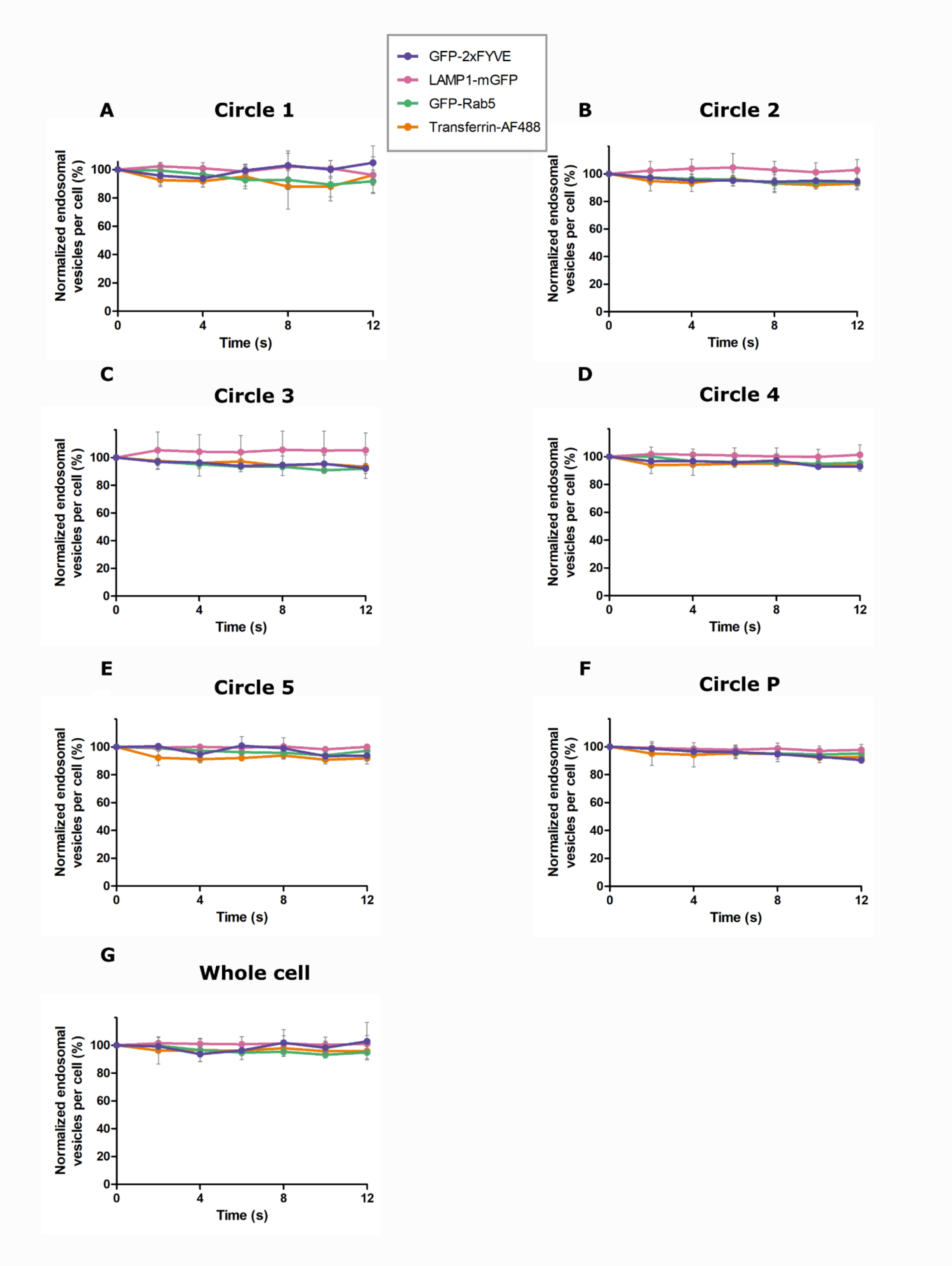


**Figure S6. Endosomal disappearance is specific to wounded cells. (A - G)** HUVEC expressing GFP-2xFYVE (purple), GFP-Rab5 (green), LAMP1-mGFP (pink), or pulsed with transferrin-AF488 for 5 min (orange), were subjected to laser ablation at a low laser power, serving as a non-wounding control. No injury to the membrane (as tracked with FM4-64 dynamics) was observable in these cases. Time-lapse images were recorded, and the endosomal disappearance was quantified as in Figure 1B. Graphs show the endosomal count across different regions of the cell from the wound site – circle 1 (A), circle 2 (B), circle 3 (C), circle 4 (D), circle 5 (E), peripheral cell edge, circle P (F), and the entire cell (G), expressed as a percentage over time (see also Figures 1C - 1H). Mean $\pm$ SD, 18 - 20 cells per marker pooled from three independent experiments.

No disappearance of the various endosomal markers is observed under these conditions indicating that the endosomal disappearance is a wounding-specific response. Multiple comparisons were performed across the endosomal proteins with the following statistical analyses: one-way ANOVA including Friedman test with $P = 0.0672$ (A), $P = 0.0600$ (B), 0.1667 (C), 0.1411 (G). For (D), (E) and (F), one-way ANOVA with Kruskal-Wallis test was performed with $P = 0.1229$ (D), $P = 0.1141$ (E) and $P = 0.596$ (F).

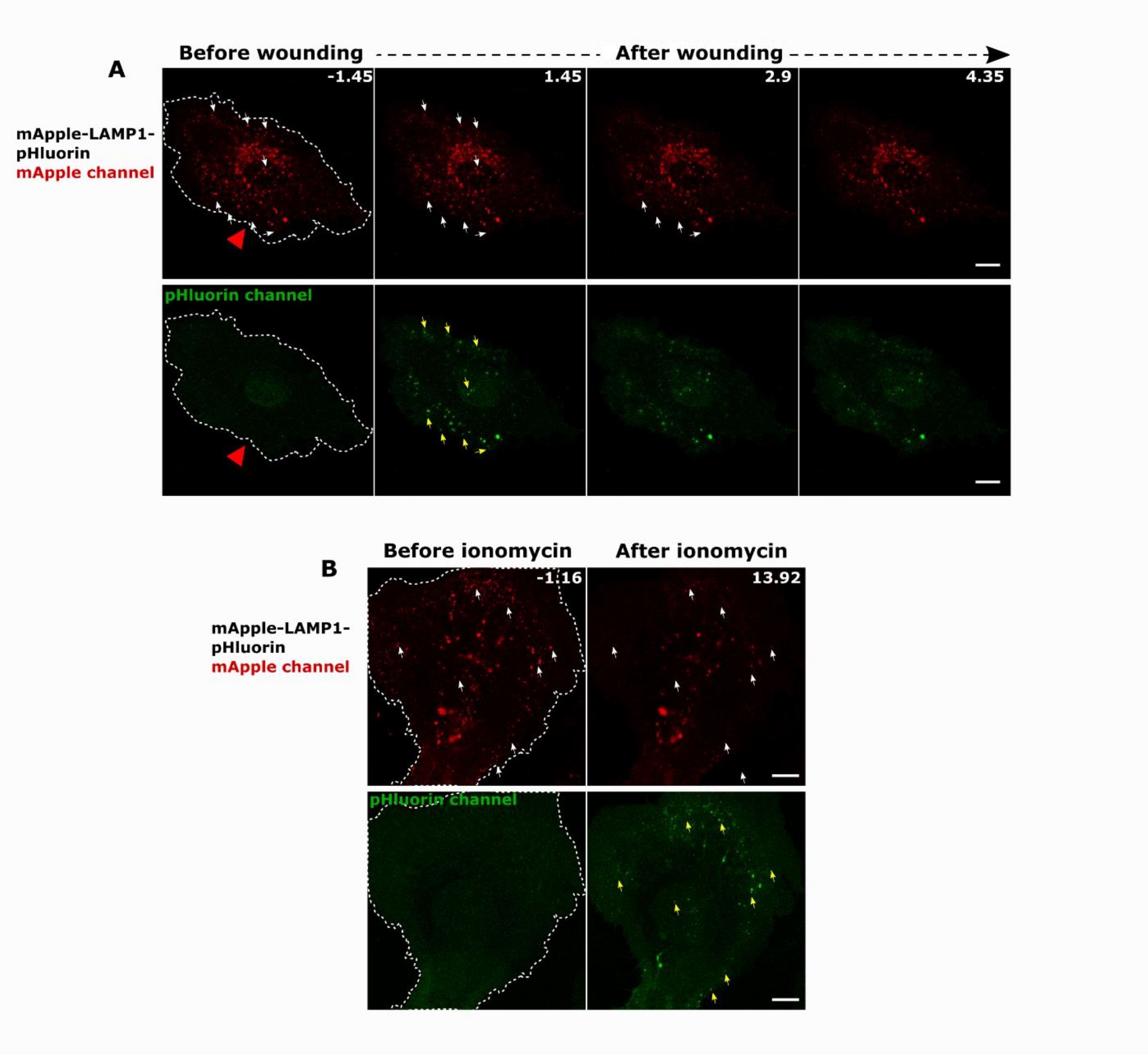


**Figure S7. Disappearance of LEL after membrane wounding is due to exocytosis. (A)** pHluorin-LAMP1-mApple was ectopically expressed in HUVEC and laser injury was performed at the lateral membrane edge (wound ROI, red triangle). Time-lapse images of the mApple channel (displayed in red) showed the disappearance of LEL as observed in Figure 1A. Images of the same cell in the pHluorin channel (green) showed an increase of fluorescence intensity after wounding at the same sites of mApple disappearance. (**B**) HUVEC transfected with the pHluorin-LAMP1-mApple construct were imaged live, ionomycin (2 µM) was added and time-lapse videos were recorded. Representative stills before and after the addition of ionomycin are shown. Again, mApple showed a disappearance accompanied by an increase in pHluorin intensity at the same sites indicating lysosomal exocytosis events. In (A) and (B), white arrows indicate LAMP1-mApple vesicles that disappear and yellow arrows indicate the corresponding exocytosis event in the pHluorin channel. White dashes, wounded (A) and stimulated (B) cells. Scale bars, 10 µm.

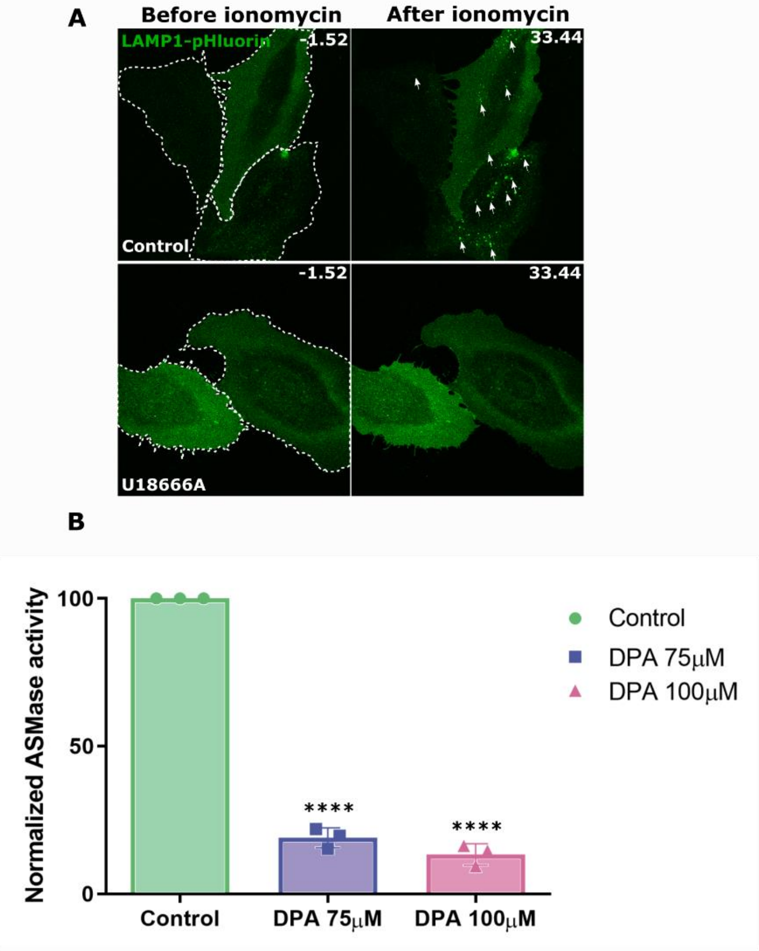


**Figure S8. U18666A is an effective inhibitor of LEL exocytosis and desipramine blocks ASM enzyme activity. (A)** HUVEC transfected with pHluorin-LAMP1-mApple were treated with U18666A (2 µg/ml) or vehicle control for 18-24 hours. Time-lapse imaging was commenced followed by the addition of 2 µM ionomycin. The pHluorin channel (green) was used to detect exocytosis events (marked with white arrows), as shown in the control. U18666A inhibited LEL exocytosis as revealed by the lack of an ionomycin-induced increase in the pHluorin signals. White dashes outline the cells prior to ionomycin stimulation. See also Figure 2C for quantification. Scale bars, 10 µm. (**B**) HUVEC were treated with desipramine - 75 µM or 100 µM, or vehicle control, for 1 h at 37°C, washed, and lysates prepared by freeze-thawing and sonication. ASM enzyme activity in the lysates was measured as described in the Methods. The ASM activity normalized to the control is represented as mean with SD from 3 independent experiments; ****$P < 0.0001$ (one-way ANOVA with Tukey’s test for comparison).

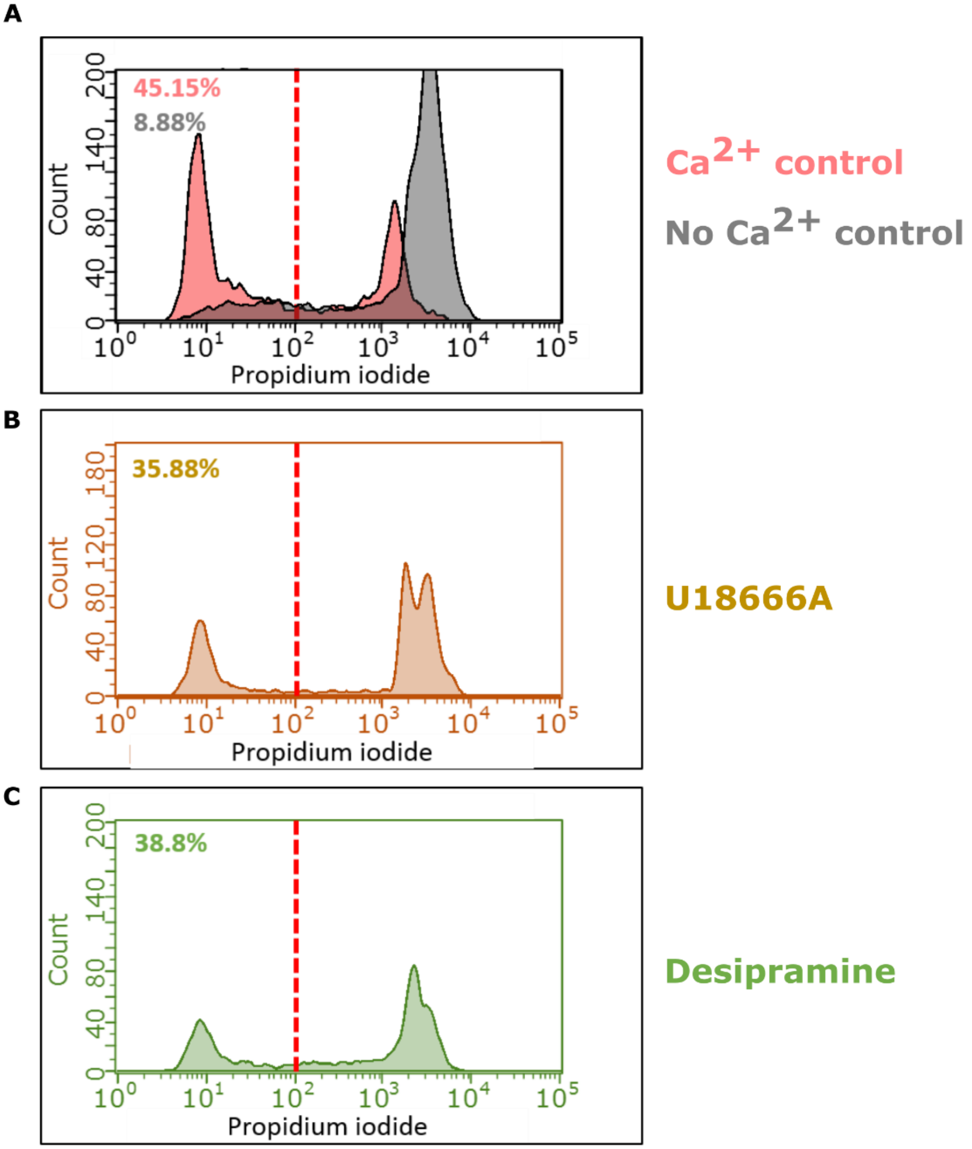


**Figure S9. Inhibition of LEL exocytosis or ASM activity does not affect HUVEC resealing.** **(A - B)** Flow cytometry profiles of HUVEC subjected to scrape injury in the presence of $Ca^{2+}$ (red, A), without $Ca^{2+}$ (grey, A) or with $Ca^{2+}$ and U18666A treatment (2 µg/ml for 20-24 hours; brown, B). Cell culture media also contained Dextran-AF488 to detect the scrape-injured cells. After incubation at 37°C for 5 min, propidium iodide (PI) was added to label the injured and non-resealed cells. The graph shown represents data from injured cells (Dextran-AF488 positive). The red dashed line indicates the gate used to calculate the population of injured and repaired cells (left offset of the dashed line). Note the increase in the population of PI-positive cells in the sample without $Ca^{2+}$ (grey), as opposed to the $Ca^{2+}$ control (red), and in the U18666A-treated sample (brown). The percentages of repaired cells are indicated in the corresponding colours on the top left of the graph. (**C**) Similar flow cytometry analysis of HUVEC scrape injured following DPA treatment (75 µM for 1 hour; marked green in the graph). Note that the cells treated with inhibitors suffered cell detachment and thereby loss of cells.

Thus, measurements were pooled from multiple technical replicates for each biological sample to ensure comparable cell numbers across conditions. All the graphs show a representative FACS profile from three independent experiments. See also Figure 2H for quantification of the flow cytometry analysis.

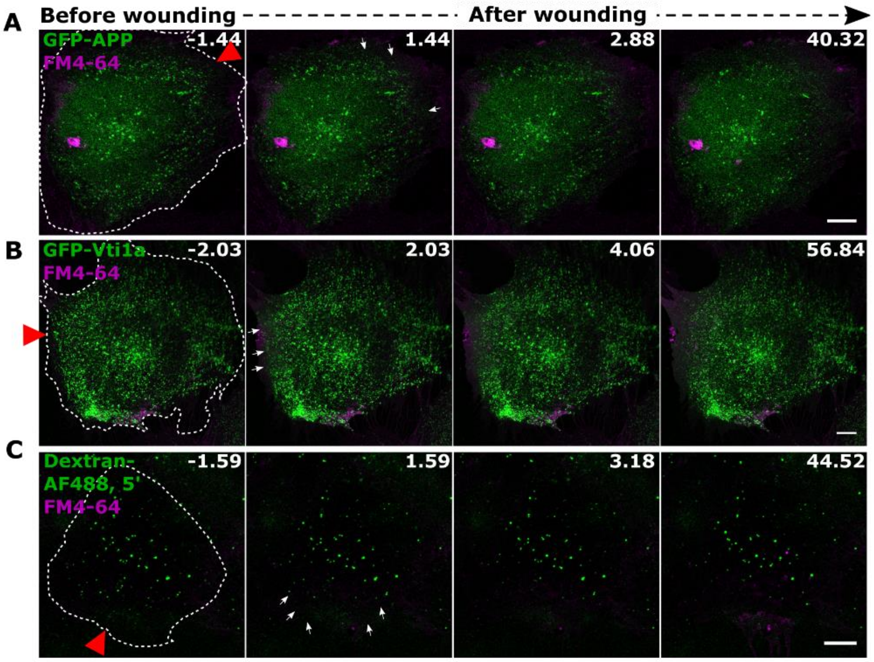


**Figure S10. Response of early endosome associated proteins to PM wounding in HUVEC. (A - B)** HUVEC transfected with various early endosome associated markers, GFP-APP (A) and GFP-Vti1a (B), were wounded by laser injury in the presence of FM4-64 (magenta and all others in green). All markers showed a disappearance near the wound site similar to GFP-2xFYVE (seen in Figure 1A). (**C**) HUVEC were pulsed with Dextran-AF488 (10 KDa) for 5 min (additional marker for freshly endocytosed vesicles; displayed in green), washed and laser injured immediately. A disappearance of early endosomes marked by dextran was also observed near the wound site. Red triangle indicates the wound site and white dashes outline the wounded cells. White arrows indicate the disappearing endosomes. Scale bars, 10 µm.

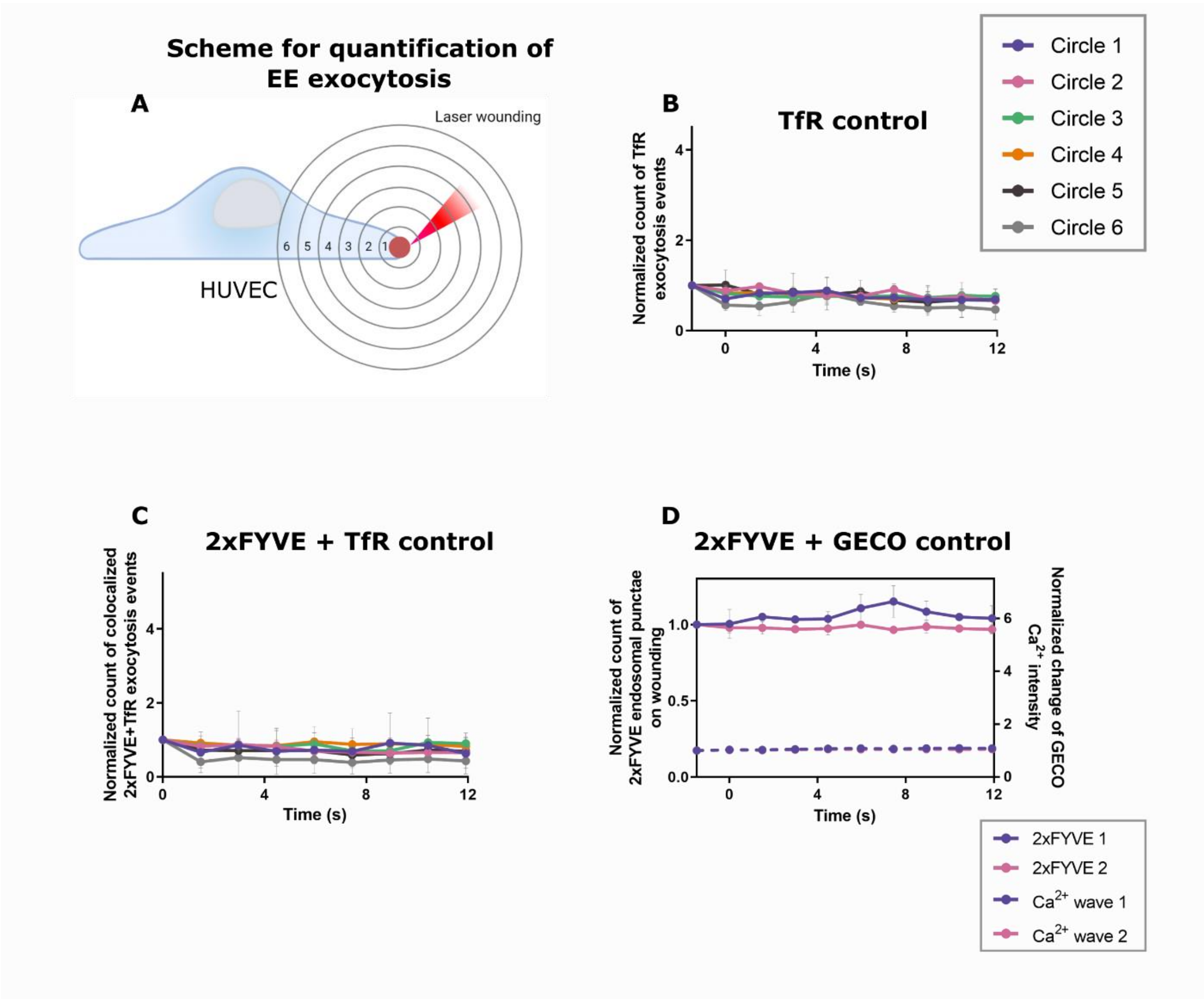


**Figure S11. Quantification of EE exocytosis with respect to the site of injury shows no response in non-wounded cells. (A)** Schematic depicting the concentric circle ROI-analysis used to quantify the early endosomal exocytosis events. The wounded cell was divided into 6 circles with increasing distance from the wound site (marked in red). Each circle was 20 µm in diameter and the measurements were always normalized to the ROI area. This ensured that the punctae count per ROI is not biased by the increasing area in the further circles. (**B – C)** Quantification of exocytosis events in non-wounded control cells showing the exocytosis count of TfR (B) and colocalized punctae of 2xFYVE disappearance and TfR exocytosis (C), following a low laser power ablation. After normalization, the events were plotted over time as for the corresponding wounded cells (see Figures 3B and 3C). Note that TfR and colocalized events of 2xFYVE disappearance and TfR punctae (2xFYVE + TfR) hardly show any change in intensity in resting cells without wounding. **(D)** Quantification of 2xFYVE-endosomal punctae disappearance and R-GECO $Ca^{2+}$ intensity change (shown in dashed lines) was carried out as in Figure 4C for control low-laser power treated cells. The disappearance of 2xFYVE punctae is plotted along the left Y-axis and the R-GECO $Ca^{2+}$ intensity along the right Y-axis, as a function of time. No evident disappearance of endosomes (as seen in Figure 1H and Figure S6A) and discernible changes in $Ca^{2+}$ intensity were noted for cells without membrane injury. $n$ = 22 cells (B and C), and $n$ = 19 (E), pooled from 3 independent experiments. For (B - C), multiple comparisons after wounding were performed with one-way ANOVA with Kruskal-Wallis test with the following $P = 0.1001$ (B), and 0.2376 (C). For (D), $P = 0.7562$ between 2xFYVE and GECO with two-tailed Mann-Whitney $U$ test was performed.

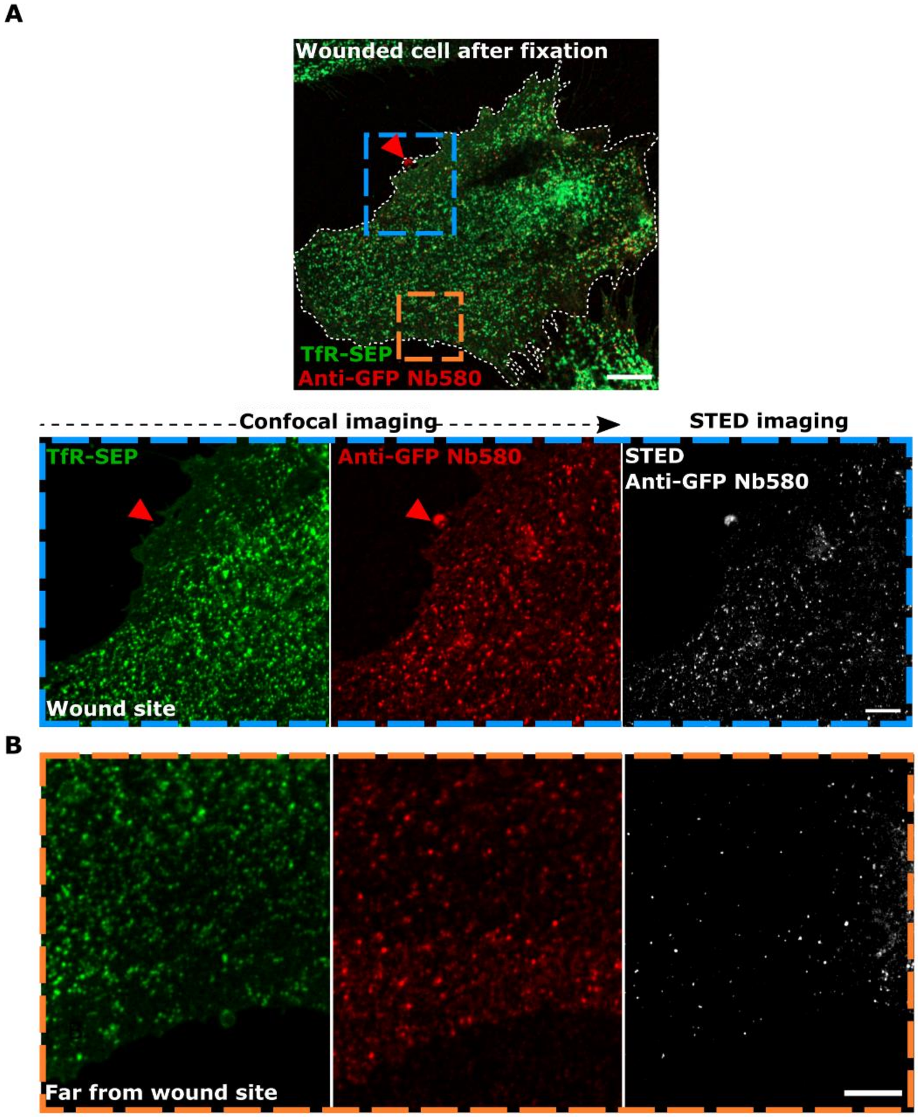


**Figure S12. Exocytosed transferrin receptors accumulate at and around the wound site. (A)** HUVEC transfected with TfR-SEP were incubated with anti-GFP nanobodies (Nb) to block the surface pool of receptors, followed by laser wounding in the presence of labelled anti-GFP Nb 580 nanobodies to stain the freshly exocytosed TfR-SEP molecules. Cells were then fixed immediately and imaged by confocal and STED microscopy. Confocal image shows a wounded cell after fixation. Dashed blue region shows the wound site magnified and an enrichment/accumulation of exocytosed TfR at and near the wound site in the Nb channel. STED image clearly resolves TfR accumulations at and near the wound site observed by the Nb clusters. Red triangle indicates the wound site and white dashes outline the wounded cell. (**B**) Imaging of a region far away from the wound site as magnified in the dashed orange box. No specific cell surface accumulation of TfR-SEP was discernible by anti-GFP Nb 580 labeling in the confocal or STED images, only background staining was observed (note the difference in anti-GFP Nb580 signal between A and B). Scale bars, 10 µm; for zoom, 5 µm.

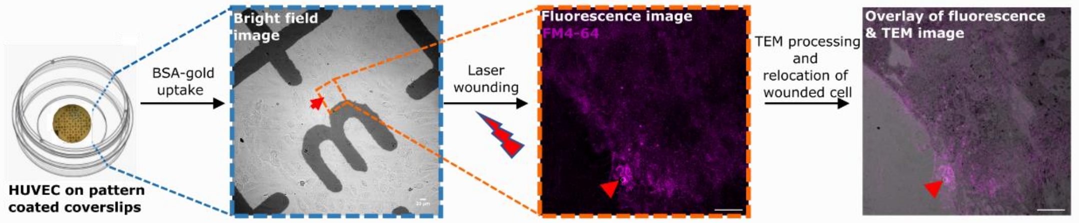


**Figure S13. CLEM analysis of the ultrastructure of a HUVEC wound site.** Schematic outline of the protocol used for imaging the wounded cell by correlative light and electron microscopy (CLEM). HUVEC were seeded on custom-made pattern-coated coverslips and a cell next to an identifiable pattern was selected (20x bright field imaging) for laser injury in the presence of FM4-64 (magenta) to mark the wound site (63x, confocal imaging). Samples were fixed 40 s after wounding and processed for TEM. The grid pattern was used to relocate the single wounded cell on the coverslip and the overlay represented shows successful CLEM relocation. Red triangle indicates wound ROI. Scale bar for bright field image, 20 µm; for other images, 10 µm.

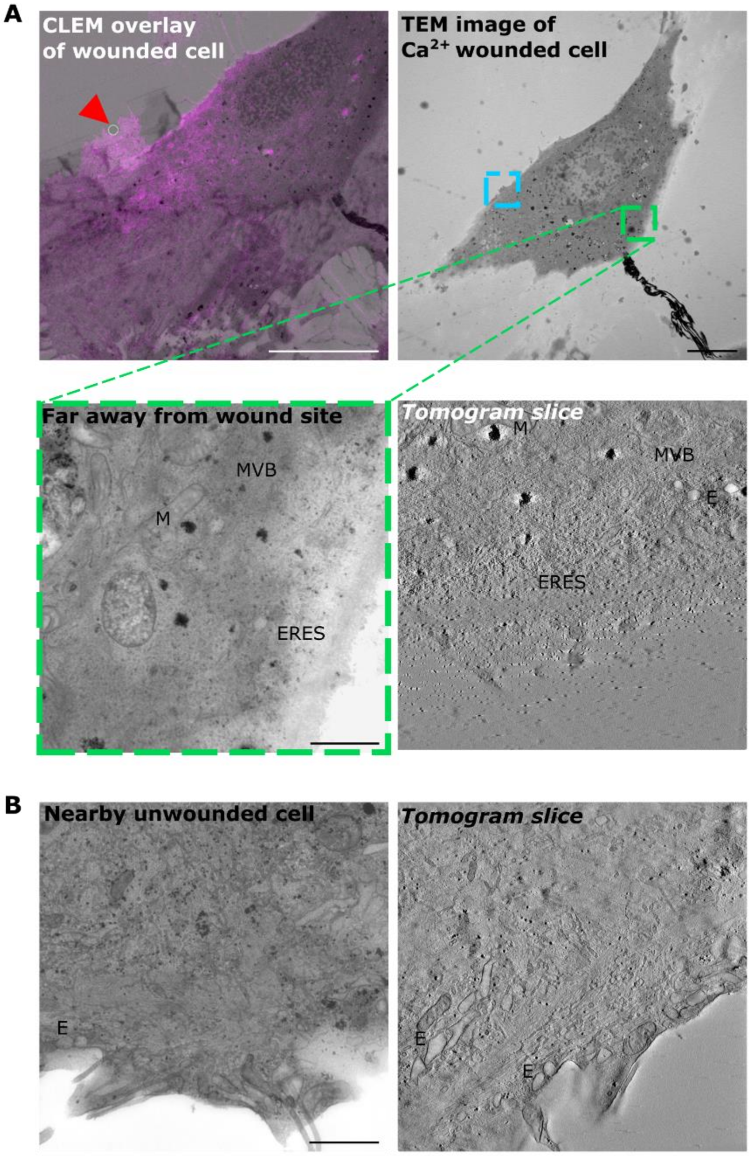


**Figure S14. CLEM imaging reveals no accumulation of vesicular structures far away from the wound site or in unwounded cells. (A)** Overlay of fluorescence and TEM images of the wounded cell (same as in Figure 3E, with wound ROI indicated by red triangle) and the corresponding TEM image. Green box indicates a region defined as far away from the wound site with the wound site marked as a blue box. Zoomed TEM image of the region far away from the wound site showed no changes in the subcortical membrane organization or morphology, as also revealed by the presence of different intracellular compartments (mitochondria indicated with M, multivesicular bodies indicated with MVB, ER exit sites indicated with ERES, endosomes indicated with E). A tomogram slice recorded at 8000x, of the same region is also displayed (for the tomogram tilts, see Video S8). Scale bars, 10 µm; for zoom, 1 µm. (**B**) TEM image of an unwounded cell as an additional control displaying no changes in the sub-membrane organization as seen for endosomes marked on the image. A tomogram slice of the unwounded cell is also shown (for the tomogram tilts, see Video S9). Endosomes indicated with E. Scale bars, 1 μm. Representative CLEM image from *n* = 17 cells (A) and *n* = 3 cells (B) across 3 independent experiments.

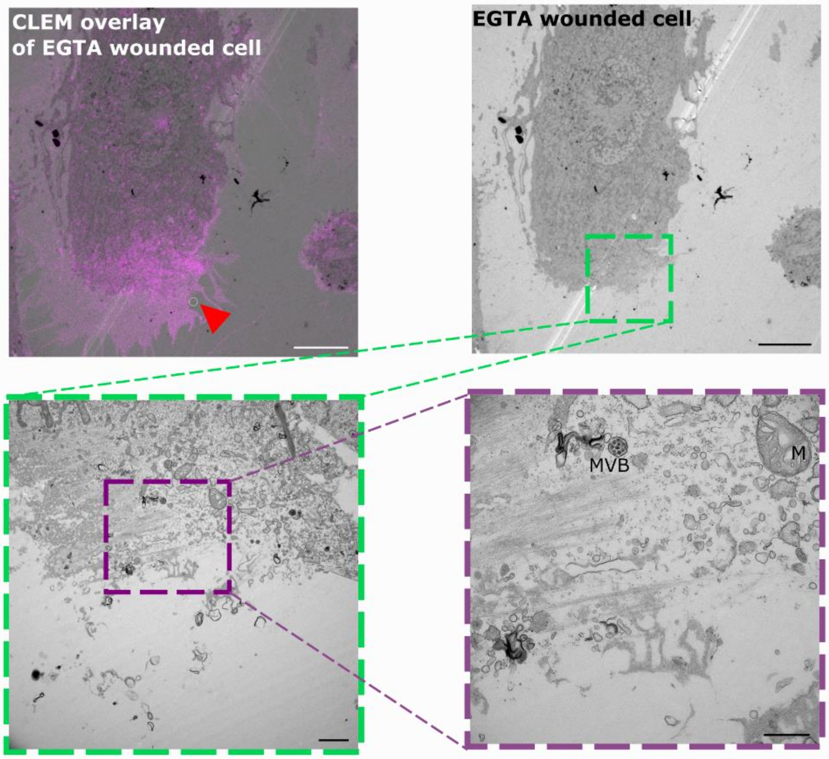


**Figure S15. HUVEC wounded in the absence of extracellular $Ca^{2+}$ show a disrupted PM and intracellular leakage.** Overlay of fluorescence and TEM image of a cell wounded in the presence of EGTA (wound ROI indicated by red triangle) as well as the corresponding TEM image of a 60 nm ultrathin section. Zoom-in of the wound site, as marked by the green box, showed massive cell damage indicating the lack of proper resealing (fragmented plasma membrane and leakage of cytoplasm). Further zoom-in, as displayed with the purple box, showed destroyed cellular organization. Mitochondria indicated with M, multivesicular bodies indicated with MVB. Scale bars, 10 µm; green box, 1 µm; purple box, 500 nm. CLEM image representative of *n* = 3 cells pooled over 3 independent experiments.

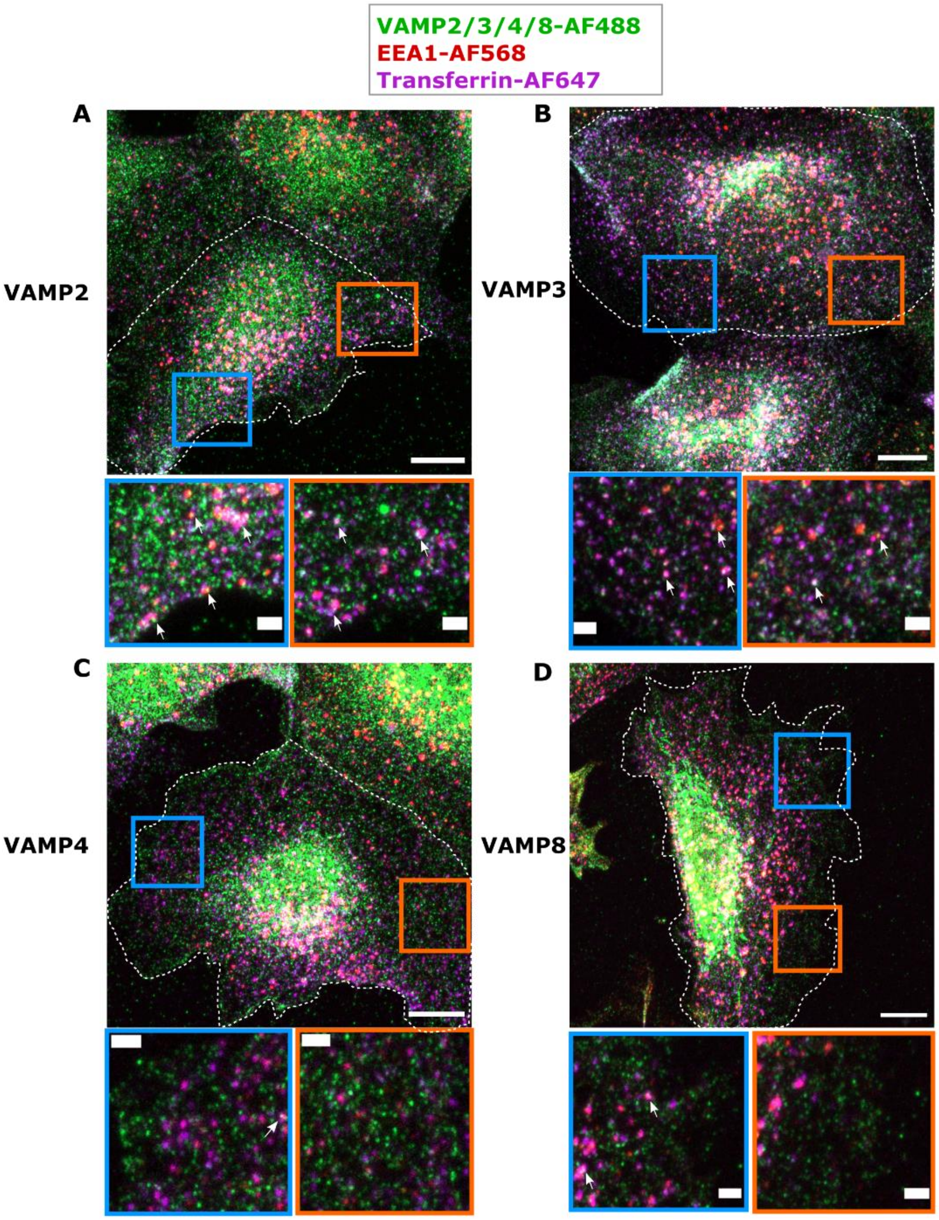


**Figure S16. Presence of early endosome associated SNARES in HUVEC. (A - D)** HUVEC were pulsed with transferrin-AF647 for 5 min to populate early endosomes, fixed and immunostained for the following VAMP proteins, VAMP2 (A), VAMP3 (B), VAMP4 (C), or VAMP8 (D), and an additional early endosomal marker, EEA1. Maximum intensity projections of the various VAMP immunostainings with the early endosomal markers are represented. Zoomed insets (marked on the whole cell) show examples of colocalization of the corresponding VAMPs with EEA1 and transferrin, as indicated by the white arrows. Cells selected for insets are outlined in white dashes. Scale bars, 10 µm; for zooms, 2 µm. The legend is given at the top of the figure in a black box. Images representative of *n* = 25-30 cells from 3 independent experiments. See also Figures 4F and 4G for quantification.

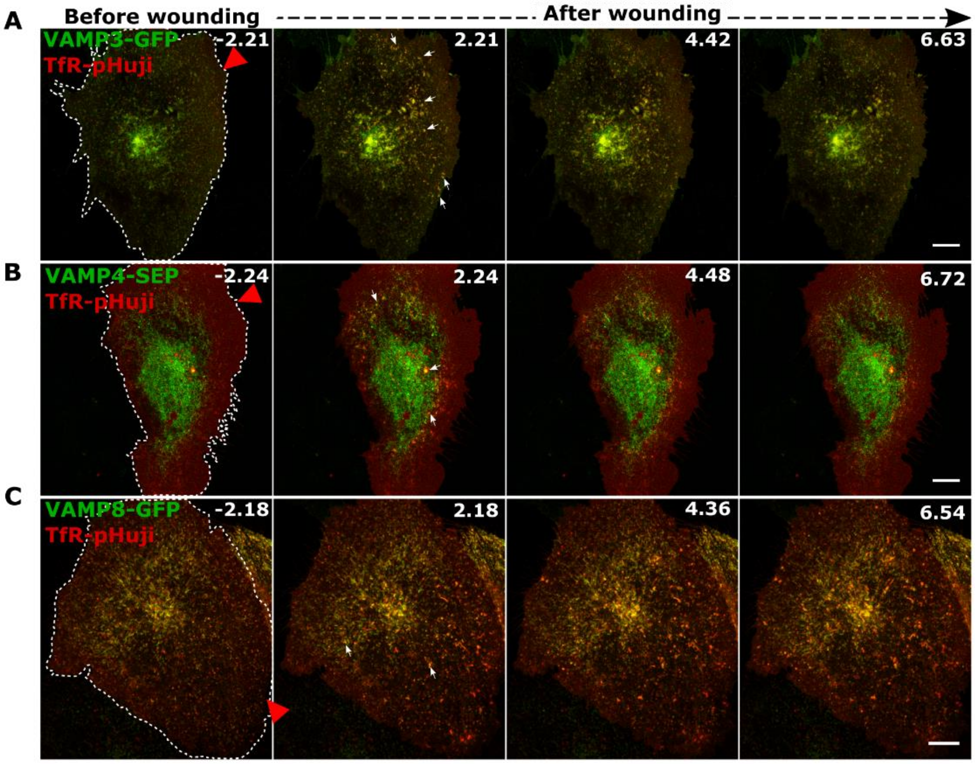


**Figure S17. Various VAMPs undergo exocytosis to different extents upon wounding in HUVEC. (A - C)** HUVEC were cotransfected with VAMP3-GFP (A), VAMP4-SEP (B), or VAMP8-GFP (C), respectively, and TfR-pHuji (A – C, pHuji displayed in red and all others in green) and subjected to laser wounding. Representative time-lapse images before and after wounding are shown. Note the relative abundance of VAMP3-positive clusters on the cell surface in panel A (increased fluorescence due to neutralization upon exocytosis), and only very few TfR-colocalized clusters of VAMP4 and VAMP8 are induced by wounding (panels B and C). Some examples of these surface clusters are marked by white arrows. Red triangle, wound ROI, and white dashes, laser injured cells. Scale bars, 10 µm.

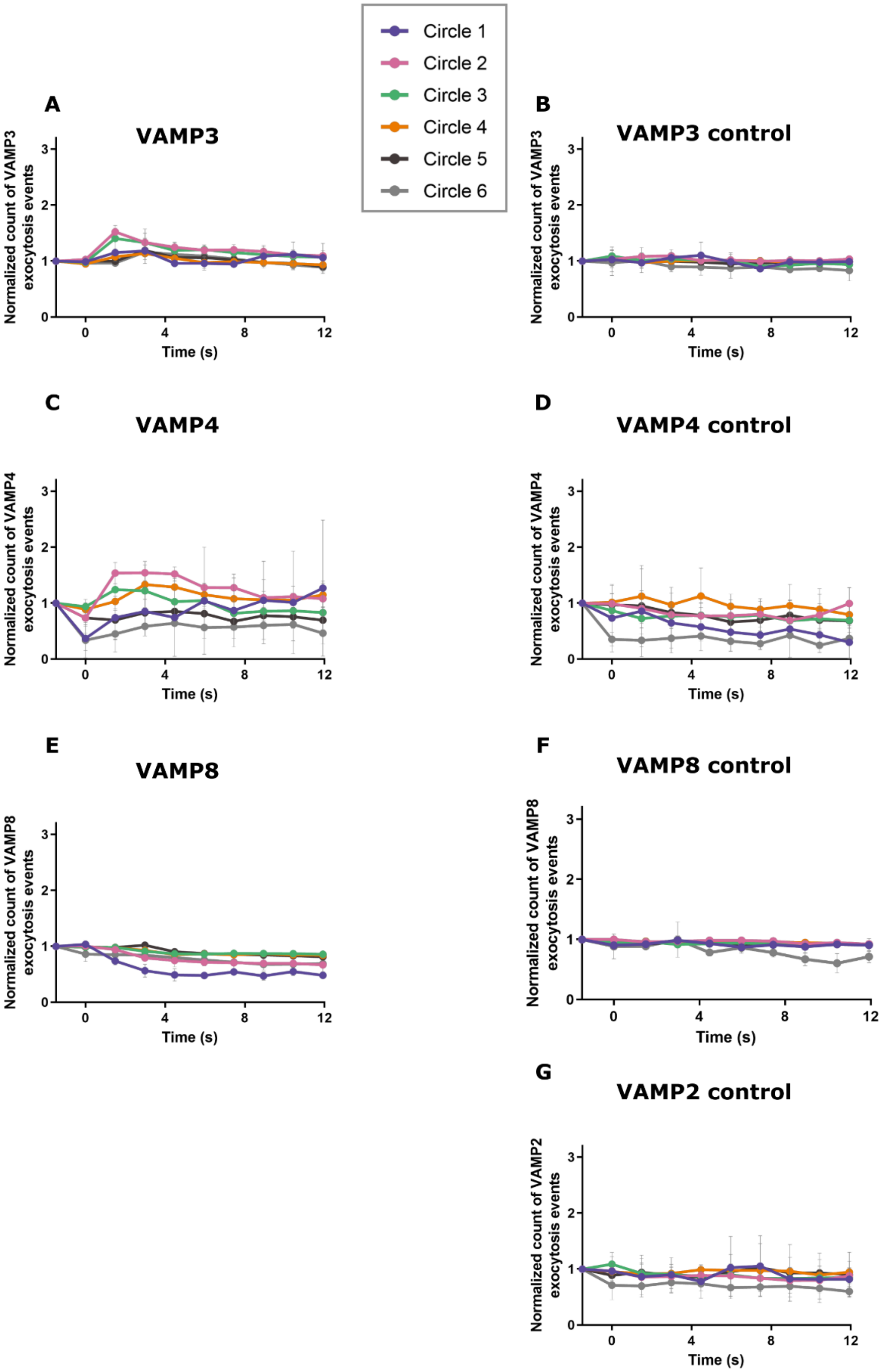


**Figure S18. Non-early endosome associated VAMPs are not presented on the cell surface upon wounding. (A - C)** Quantification of surface clusters based on increased fluorescence of

VAMP3- GFP (A), VAMP4-SEP (B), and VAMP8-GFP (C) after laser wounding (as in Figure S17). The clusters were quantified after iLastik pixel-based thresholding of images and counted across various circles from the wound ROI after injury (see Figure S11A). The values are represented as the increment in VAMP clusters normalized to the baseline count before wounding and to the area in $\mu m^2$ of each ROI. Note the minimal increase in exocytosis (i.e. fluorescence increase due to neutralization) after wounding in the various VAMPs analysed, as compared to VAMP2 (Figure 5B) indicating that not all VAMPs participate in injury mediated EE exocytosis. **(D – F)** Non-wounded control cells were subjected to similar quantification for VAMP3-GFP (D), VAMP4-SEP (E), and VAMP8-GFP (F), by laser ablation at a low laser power, whereby no membrane injury was induced. Note that the exocytosis of various VAMPs after wounding is comparable to the corresponding non-wounded control. This indicates that non-early endosome associated VAMP 3/4/8 barely undergo exocytosis following membrane injury. **(G)** Quantification of VAMP2-SEP exocytosis events in non-wounded cells quantified as above (corresponding control graph for Figure 5B). No observable exocytotic events were seen in resting VAMP2-SEP expressing cells. The legend is given at the top of the figure in a black box. Mean $\pm$ SD shown here with $n$ = 20 cells (A, C, D), $n$ = 16 (B), $n$ = 18 (E), and $n$ = 17 (F), pooled from 3 independent experiments. Statistical comparisons were performed as follows: repeated measures ANOVA with $P$ = 0.0312 (A) and $P$ = 0.0539 (B); one-way ANOVA with Kruskal-Wallis test with $P$ = 0.0576 (C), $P$ = 0.2030 (E), and $P$ = 0.3542 (G); and ordinary one-way ANOVA with $P$ = 0.8880 (D) and $P$ = 0.1875 (F).

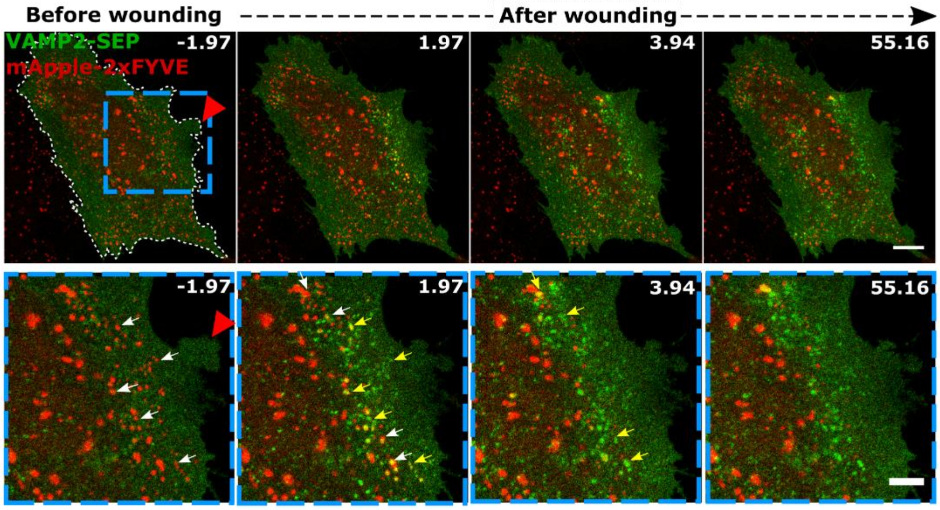


**Figure S19. VAMP2 exocytosis occurs at sites of early endosome disappearance.** HUVEC were transfected with VAMP2-SEP (green) and mApple-2xFYVE (red) and subjected to laser wounding. Representative images of a cell pre and post wounding are shown. Dashed blue box (magnified below) shows that the sites of VAMP2 exocytosis (increase in fluorescence due to neutralization after fusion with the PM) correlate with mApple-2xFYVE endosomal disappearance. White arrows indicate the 2xFYVE endosomes before disappearance. Yellow arrows indicate the colocalized sites of VAMP2 cluster formation and 2xFYVE disappearance. Red triangle, wound ROI and white dashes, laser ablated cell. Scale bars, 10 µm; for zoom, 5 µm.

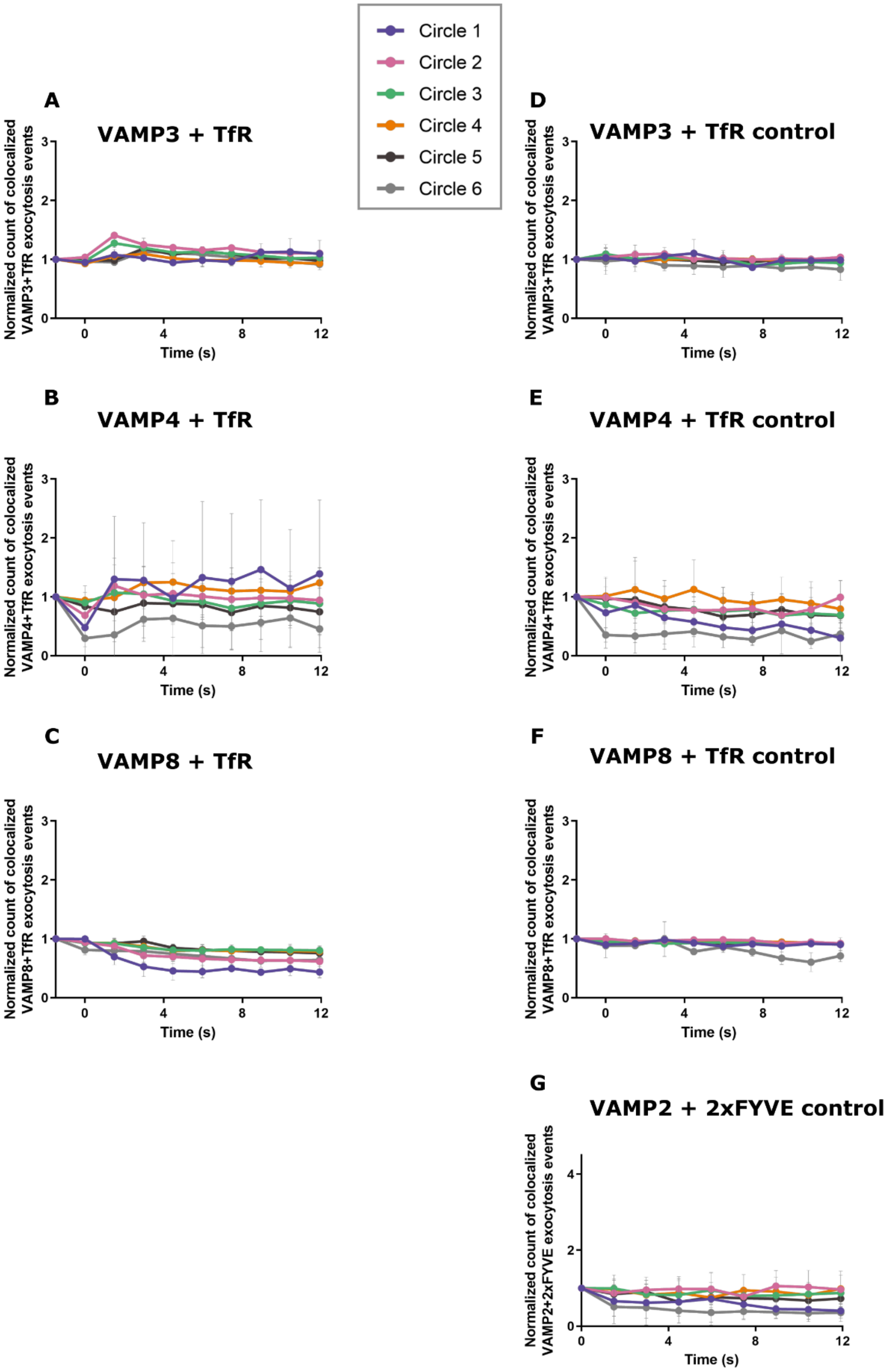
Circle 1
Circle 2
Circle 3
Circle 4
Circle 5
Circle 6
A
VAMP3 + TfR
B
VAMP4 + TfR
C
VAMP8 + TfR
D
VAMP3 + TfR control
E
VAMP4 + TfR control
F
VAMP8 + TfR control
G
VAMP2 + 2xFYVE control
Normalized count of colocalized VAMP3+TfR exocytosis events
Normalized count of colocalized VAMP4+TfR exocytosis events
Normalized count of colocalized VAMP8+TfR exocytosis events
Normalized count of colocalized VAMP2+2xFYVE exocytosis events
Time (s)

**Figure S20. VAMP 3, 4 and 8 are not associated with wounding induced EE exocytosis events. (A - C)** Colocalization of VAMP3-GFP (A), VAMP4-SEP (B), or VAMP8-GFP (C), with TfR exocytosis events upon wounding was quantified as for VAMP2 (in Figure 5C). The thresholded images for each VAMP and TfR were compared for colocalized punctae in the same frame following wounding. This was quantified across the different ROIs from the wound site (see Figure S11A). The values are normalized to the initial baseline number of random colocalized punctae before wounding and per $\mu m^2$ of each ROI. **(D – F)** Quantification of colocalized punctae of VAMP3 (D), VAMP4 (E), or VAMP8 (F), with TfR in resting non-wounded cells. Note that there are virtually no differences in the different VAMP + TfR exocytosis events between wounded and control cells. **(G)** Quantification of exocytotic events of VAMP2-SEP overlapping with GFP-2xFYVE disappearances for control non-wounded cells. Similar analysis as in Figure 5E, with images being subjected to walking average and colocalized punctae counted across ROIs over time. The legend is given at the top of the figure in a black box. Mean $\pm$ SD shown here with $n$ = 20 cells (A, C, D), $n$ = 17 (B), $n$ = 18 (E), and $n$ = 17 (F), pooled from 3 independent experiments. Multiple comparisons after wounding were performed for the above datasets using one-way ANOVA with Kruskal-Wallis test with the following $P$ values: 0.0298 (A), 0.329 (B), 0.0836 (C), 0.2030 (E), 0.2029 (F), and 0.4602 (G). For (D), ordinary one-way ANOVA was performed with $P$ = 0.3256.

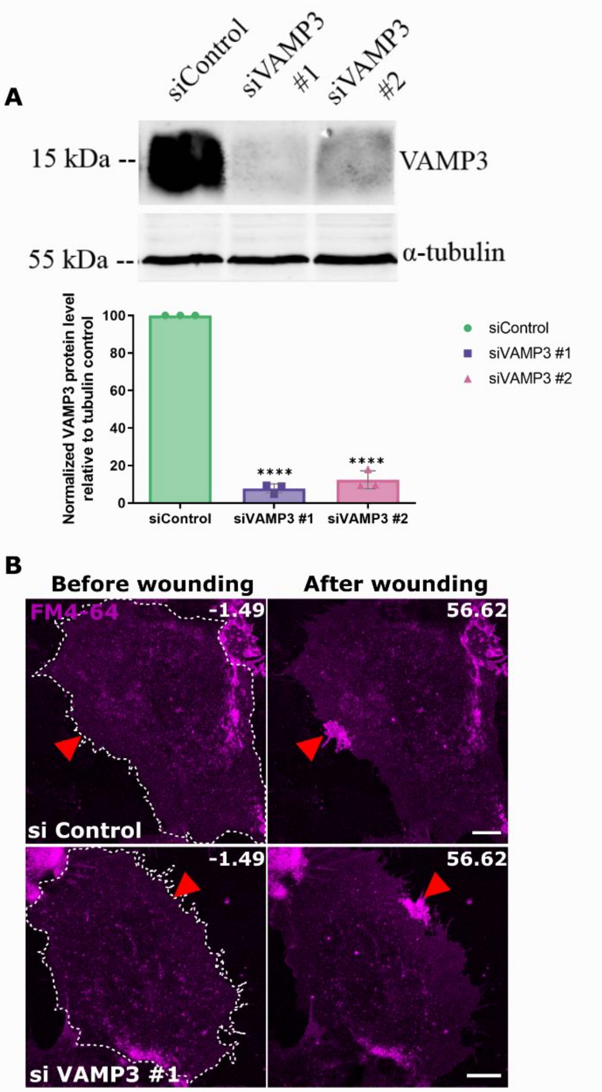


**Figure S21. VAMP3 does not contribute to HUVEC membrane resealing. (A)** Western blot showing protein levels of VAMP3 in HUVEC after siRNA transfection with siControl or two different VAMP3 siRNAs (siVAMP3 #1 and siVAMP3 #2), in the top panel. As a loading control, α – tubulin was blotted (bottom panel). The graph at the bottom shows the analysis of knockdown efficiency of VAMP3 following siRNA transfection plotted as a percentage normalized to the loading control. Significant depletion of VAMP3 levels was observed with the different siRNAs against VAMP3. A representative blot is shown and mean ± SD plotted from 3 independent experiments in the graph. ****$P < 0.0001$ (one-way ANOVA with Hol-Sidak's multiple comparison test used here). **(B)** Laser wounding of siControl or siVAMP3 (siVAMP3 #1 shown here) transfected HUVEC in the presence of FM4-64 (magenta). Representative time-lapse images before and after wounding are shown. Efficient membrane resealing was observed in siControl and all siVAMP3 transfections, indicating that VAMP3 is not functionally involved in membrane repair in HUVEC. Representative image from 4 independent experiments. Red triangle, wound site, and white dashes, wounded cells. Scale bars, 10 µm.

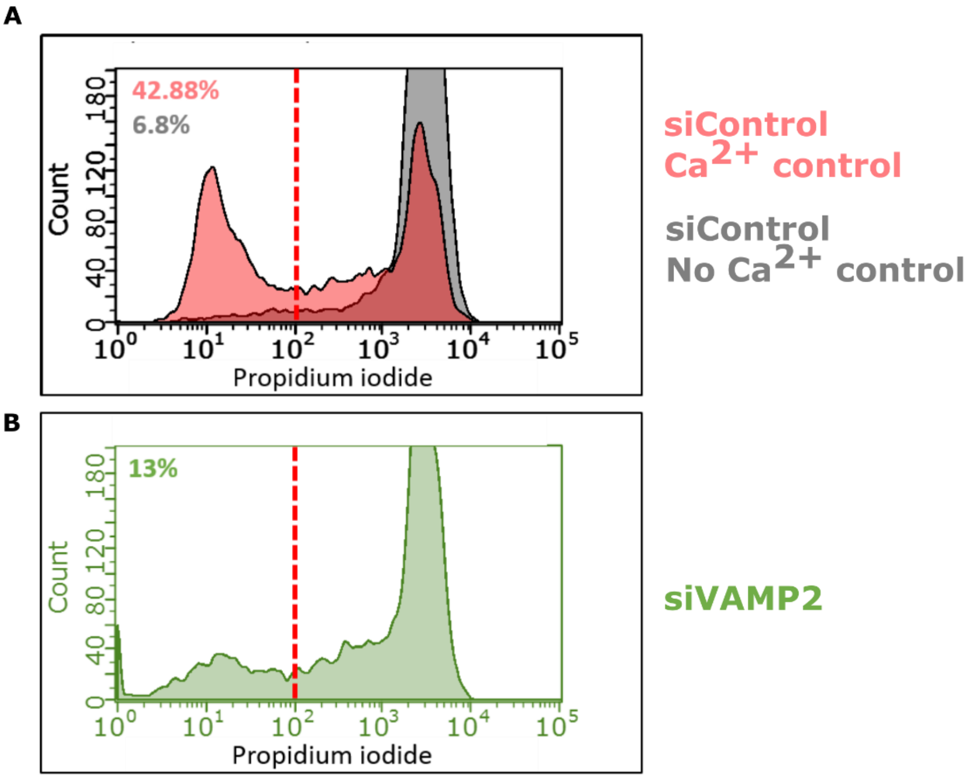


**Figure S22. Resealing defects in VAMP2 depleted cells as revealed by scrape injury assay. (A - B)** Flow cytometry profile of scrape-injured control siRNA transfected HUVEC in the presence of $Ca^{2+}$ (red, A) and without $Ca^{2+}$ (grey, A) or VAMP2 Pool siRNA transfected HUVEC in the presence of $Ca^{2+}$ (green, B). Propidium iodide staining reveals the population of non-repaired cells and the red dashed line indicates the gate used to calculate the resealed cell population ((left offset of the dashed line). The percentages of repaired cells are indicated in the corresponding colours on the top left of the graph. Note the shift in the peaks of VAMP2 siRNA treated cells to the right indicating a higher percentage of non-repaired cells (B). The graphs show a representative FACS profile from five independent experiments. See also Figure 6G for quantification of the flow cytometry analysis.

**Video S1.**

Time lapse laser ablation recording that shows the disappearance of EE vesicles (GFP-2xFYVE, green) upon wounding (FM4-64 in magenta), related to Figure 1A. Red triangle, wound ROI. White arrows indicate 2xFYVE vesicles that disappear after wounding. Frames were captured at 1.98 *s* intervals. Time is displayed in seconds and t = 0 *s* represents the time of wounding. Scale bar, 10 μm.

**Video S2.**

Time lapse laser ablation recording that shows the disappearance of LEL (LAMP1-mGFP, green) upon wounding (FM4-64 in magenta), related to Figure 1A. Red triangle, wound ROI. White arrows show the disappearing LAMP1 vesicles after wounding. Frames were captured at 2 *s* intervals. Scale bar, 10 μm.

**Video S3.**

Time lapse laser ablation recording that shows the disappearance of EE (Transferrin-AF488, green) upon wounding (FM4-64 in magenta), and corresponds to the time series in Figure S3G. Red triangle, wound ROI. White arrows indicate the disappearance of transferrin-loaded vesicles after wounding. Frames were captured at 1.98 *s* intervals. Scale bar, 10 μm.

**Video S4.**

Time lapse laser ablation recording that shows the disappearance of EE (GFP-2xFYVE, green) and the concomitant appearance of transferrin receptor (TfR-pHuji, red) accumulations after membrane wounding, related to Figure 3A. Red triangle, wound ROI. White arrows indicate a few examples of the formation of surface TfR cluster at sites of 2xFYVE disappearance post wounding. Images were taken every 1.49 *s*. Scale bar, 10 μm.

**Video S5.**

Time lapse recording that shows the localization of EE vesicles (GFP-2xFYVE, green) after stimulation of an intracellular $Ca^{2+}$ rise by 250 nM histamine and corresponds to the images in Figure 4D. White arrows exemplify 2xFYVE vesicles that do not undergo disappearance following histamine stimulation. Note that the vesicles are very dynamic and move around in the images and in *z*-direction, which is distinguished from a complete disappearance by their reappearance in the plane of focus in later frames. t = 0 *s* represents the time of histamine addition. Frames were captured at 1.47 *s* intervals. Video shows the recording after the focus shift was corrected, as indicated in the timestamps. Scale bar, 10 μm.

**Video S6.**

Double tilt tomogram recording of a 250 nm thick section of the resealed wound site of HUVEC (from 250 – 500 nm height) at 12000x magnification, corresponding to the blue box in Figure 3E. The video scrolls through the tomographic volume in Z and the slices were then contoured for the structures as in Figure 3E. The 3D model view is displayed at the end. PM (in magenta) and endosomes of vesicular (dark blue) and tubular (light blue) shapes are traced.

**Video S7.**

Double tilt tomogram recording of a 250 nm thick section of a region nearby the wound site (from 250 – 500 nm height) at 12000x magnification, corresponding to the orange box in Figure 3E. The video scrolls through the tomographic volume in Z and the slices were then contoured for the structures as in Figure 3E. The 3D model view is displayed at the end. PM (in magenta) and vesicular endosomes (dark blue) are traced.

**Video S8.**

Tilt tomogram recording of a 250 nm thick section of a region far away from the wound site (from 250 – 500 nm height) at 8000x magnification, corresponding to the green box in Figure S14A.

**Video S9.**

Tilt tomogram recording of a 250 nm thick section of a region of a nearby unwounded cell (from 250 – 500 nm height) at 8000x magnification corresponding to the tomogram slice in Figure S14B.

**Video S10.**

Time lapse laser ablation recording that shows the appearance of VAMP2-SEP (green) and the transferrin receptor (TfR-pHuji, red) accumulations after membrane wounding, related to Figure 5A. Red triangle, wound ROI. White arrows indicate the colocalized sites of VAMP2 and TfR accumulations post wounding. Frames were captured at 1.49 *s* intervals. Scale bar, 10 μm.
